# Deletion of Nrf1 exacerbates oxidative stress-induced cellular senescence by disrupting the cell homeostasis

**DOI:** 10.1101/2024.03.09.584196

**Authors:** Da Lyu, Meng Wang, Lu Qiu, Shaofan Hu, Yiguo Zhang

## Abstract

Cellular senescence has been accepted as a fundamental contributor to ageing and a variety of age-related diseases, in which oxidative stress has been further recognized to play a critical initiation role. However, the anti-senescence potential of antioxidant nuclear factor erythroid-derived 2-like 1 (Nrf1, encoded by *Nfe2l1*) remains elusive to date, even though the hitherto accumulating evidence demonstrates that it is an indispensable redox-determining transcription factor for maintaining cellular homeostasis and organ integrity. Herein, we discovered that deletion of Nrf1 resulted in markedly elevated senescence characteristics in *Nrf1α*^−/−^ cells, as characterized by two distinct experimental models induced by oxidative stress, which are evinced by typically heightened activity of senescence-associated β-galactosidase and progressive senescence-associated secretory phenotype (SASP), along with decreased cell vitality and intensified cell cycle arrest. Further experimental investigation also uncovered that such acceleration of oxidative stress-induced senescence resulted from heightened disturbance in the cellular homeostasis, because deficiency of Nrf1α leads to the STAG2- and SMC3-dependent chromosomal stability disruption and autophagy dysfunction, though as accompanied by excessive accumulation of Nrf2 (encoded by *Nfe2l2*). The aberrant hyperactive Nrf2 cannot effectively counteract the escalating disturbance of cellular homeostasis caused by *Nrf1α*^−/−^. Overall, this study has provided a series of evidence supporting that Nrf1 indeed exerts an essential protective function against oxidative stress-induced cellular senescence, thereby, highlighting its primary indispensable contribution to maintaining robust cell homeostasis.

## 1. Introduction

Ageing rises by a multifactorial, multifaceted, progressive degenerative decline in the physio-pathological process that affects normal functional and structural integrity of higher organisms, and thereby poses a severe threat to the healthy survival and even increases the risk of degenerative diseases in the morbidity and mortality [1–3]. Such a global ageing phenomenon has been continuing and further accelerating to date [4], leading to a high rise in the prevalence of morbidity due to age-related illnesses (e.g., heart failure, diabetes, atherosclerotic disorders, neurodegenerative diseases and cancer). Currently, this trend presents significant challenges in both domains of medicine and socio-economics [5, 6]. To address this issue, it is of crucial importance to explore the endogenous potential for increased longevity and devise superior anti-ageing interventions. And many studies on the relevant mechanisms have highlighted certain markers or pillars of ageing, such as genomic instability, progenitor exhaustion/dysfunction, and cellular senescence and dyshomeostasis [7–9]. Amongst these factors, cellular senescence has directly given rise to significantly ageing process and various age-associated disorders. This underscores the potentiality of targeting senescent cells to be paved as an effective tactics for strategically extending the healthy lifespans, and also preventing and treating age-related chronic diseases [10–12]. Cell senescence is accepted as a response to acute or chronic injuries stemming from intrinsic stress and extrinsic stimuli [7]. Of striking note, free radicals and relevant reactive species (including reactive oxygen species (ROS)) are significantly involved in the initiation and progression of cellular senescence and ageing-associated diseases, because they are functioning, not only as harmful molecules, but also as signaling molecules [13]. The impact of ROS varies depending on its physio-pathological concentrations that are allowed for a dual distinctive and even opposite role of each reactive species in these processes. For example, the mild levels of ROS can enhance the defense response by inducing endogenous activation of adaptive antioxidant systems and relevant detoxifying reactions, thereby promoting the eustress-defensing tolerance and longevity. Yet, an overabundance of oxidative distress stemming from the heightened levels of ROS, that exceed the threshold of cellular tolerance and/or cause an impairment of antioxidant mechanisms within the cell, can lead to severe oxidative damages to DNA, lipids, and proteins, ultimately triggering the initiation of cell senescence and ensuing advancement [14]. With increasing ages, these collective damages are further cumulated by chronic overstimulation of oxidative stress arising from an imbalance between ROS and antioxidant defense systems and/or disrupted redox signaling control. Such gradually cumulative oxidative stress-induced damages have been accepted as ‘a redox theory of ageing’ [15], because they indeed play a crucial role in accelerating cellular senescence and even organismal aging, as bad as contribute to development of age-related disorders.

Amongst antioxidant transcription factors of the cap’n’collar-basic-region leucine zipper (CNC-bZIP) family [16], nuclear factor erythroid-derived 2-like 1 (Nrf1, encoded by *nfe2l1*) and 2 (Nrf2, encoded by *nfe2l2*) are two highly-conserved principle members expressed widely in various tissues and cells of vertebrates. Of sharp note, the living fissile-like Nrf1 is rather much closer than the water-soluble Nrf2 to their commonly-shared ancestral orthologues SKN-1, CncC and Nach proteins emerged from the early evolutionary stages of life histories [17]. The important roles of Nrf2 and even SKN-1 in oxidative stress response to ageing have been well documented [18, 19], but it is heretofore unknown about whether and/or how Nrf1 fulfills its essential cytoprotective functions in the oxidative stress-induced ageing [20].

In the normal physiological circumstances, Nrf1 is topologically localized in the endoplasmic reticulum (ER) and its ER luminal-resident domains are hence protected by this organelle membranes [21], even though its cytoplasmically-residing domains are likely to experience proteolytic degradation by predominant ubiquitin-proteasome system [22, 23]. Upon stimulation by oxidative ER stress, Nrf1 is then subjected to a sequence of topovectorial processes, involving its retro-translocation across ER membranes, selective proteolytic cleavage by DDI1/2 or other cytosolic proteases, specific modification and ensuing dislocation into the nucleus in order to regulate a unique set of target genes [24, 25]. Of importance, Nrf1 is identified to serve as a pivotal regulator in upholding the constitutive expression of intrinsic stress-defense genes governed by antioxidant response elements (AREs) through the formation of its functional heterodimers with each of sMaf or other bZIP proteins, thereby guaranteeing the maintenance of redox homeostatic balance [26–28]. The accumulating evidence has suggested that impaired Nrf1 function is much more likely to trigger the occurrence and progression of ageing-related pathologies (e.g., cardiovascular diseases, inflammatory diseases, neurodegenerative diseases), which are evidenced to be closely associated with compromised adaptive responses (i.e., both responses to oxidative and proteotoxic stresses monitored by Nrf1). Conversely, either pharmacological or phytochemical agents that promote Nrf1 activation demonstrated a considerable promise to act as potential targets for the prevention or treatment of age-related disorders [20, 29]. Thereby, it is inferable that, as a key regulator of the antioxidant system, Nrf1 has shown its significant potential in protecting cells and organisms from cellular senescence and ageing. However, the precise functional characteristics of such advantageous anti-ageing and the underlying molecular mechanisms driving it remain elusive to date.

During the senescent process, there is a noticeable and gradual change in the functionality of a cell across its multiple levels, including cellular, subcellular, and molecular [7]. Hence, in order to gain an insight into the cytoprotective role of Nrf1 against senescence induced by oxidative stress, we have herein conducted a series of experiments to evaluate alterations in various senescence-related markers, e.g., increased activity of senescence -associated β-galactosidase (SA-β-gal) and specific expression of senescence-associated secretory phenotype (SASP), along with the cell growth arrest, in two distinct model cell lines (HepG2 and HeLa) with the indicated full-length Nrf1α-specific knockout (*Nrf1α*^−/−^) within established models of oxidative stress-induced senescence. Since the protective function of Nrf1 is closely associated with its unique ability to play a beneficial role in maintaining robust cell homeostasis under normal or even stressful conditions [22, 26], the objective of this study is targeted to elucidate the molecular mechanisms underlying the homeostatic protective role Nrf1 as a prospective strategy to counteract oxidative stress-induced senescence. For this aim, we have further investigated this functional effects of Nrf1 and relevant molecular underpinnings in the cellular senescence insofar as to establish a theoretical foundation for developing a new tactic to prevent and treat cellular senescence and ageing, and also to provide an insight into potential chemotherapeutic interventions for those ageing-related diseases by precision targeting Nrf1.

## 2 Experimental procedures

### 2.1 Cell culture and chemical solvents

HepG2 cells were originally sourced from American Type Culture Collection, whilst HeLa cells were from the National Infrastructure for Cell Line Resource. Additionally, short tandem repeat (STR) typing maps and authentication profiles were also employed to confirm the fidelity of both cell lines (performed by Shanghai Biowing Applied Biotechnology Co., Ltd., Shanghai, China). Of note, both *Nrf1α*^−/−^ and *Nrf2*^−/−^ cell lines were previously generated by Qiu *et al* [30], using their gene editing of wild-type (WT) HepG2, and similar cell lines were established on the base of HeLa cells in our laboratory, all of which were subsequently identified in this study (Fig. S1). All these experimental cells were incubated at 37°C with 5% CO_2_ and cultured in basic DMEM (Gibco, Grand Island, NY, USA) supplemented with 10% of FBS (Gibco) and 1% of antibiotics (penicillin–streptomycin, Gibco). All other chemicals, reagents, software tools and equipment resources used in this work are listed in Supplementary Table 1.

### 2.2. Cell viability assayed by MTT

To analyze the viability of experimental cells that had been cultured overnight in 96-well plates (on a flat bottom) at a plating concentration of 1 × 10^2^ cells/μL, the MTT assay was performed appropriately according to experimental requirements. At the end of treatment, the cultured cells in indicated wells were appended with MTT (5 mg/mL, Molekula, Darlington, UK) and continuously maintained for 4 h at 37 ° C in the culture incubator. Subsequently, the solution within the orifice was removed, before dimethyl sulfoxide (Solarbio) was dispensed to solubilize the reaction product. The absorbance readings were taken at a wavelength of 490 nm.

### 2.3 SA-β-gal staining and its activity assay

The experimental cells were appropriately manipulated as required for an SA-β-gal (senescence-associated β-galactosidase) staining kit (CST, Danvers, MA, USA), which was employed to assess cellular senescence following the manufacturer’s protocol instruction. After indicated treatment of cells, they were washed twice by PBS and then fixed in a formulated fixative for 15 min at room temperature. The fixed cells were further washed thrice by PBS and subsequently incubated overnight in a staining solution at 37°C without additive CO_2_. The images of the blue stained cells were observed under a microscope and quantified as the β-gal positivity. The cytosolic SA-β-gal levels were further measured by another assay kit of its activity (Solarbio). All the samples and standard solutions were prepared, sonicated, and processed according to the company-provided instructions, before their optical density (OD) was determined at 400 nm. The BCA methodology was also employed to quantify total proteins extracted by using a protein assay reagent kit (ApplyGen, Beijing, China). Lastly, the β-gal activity was determined and calculated by the specific formula described in the protocol.

### 2.4 Two distinct models of cell senescence

Experimental cell lines were allowed for growth in 96-well culture plates (on the flat bottom) until reaching 80% confluence and subsequently changed in fresh culture media, before being subjected to establishment of two distinct models of cell senescence. Firstly, to establish a simplified senescence model (i.e., Model 1), the cells within a rather highly confluence were continuously cultivated for distinct periods of time (e.g., 24, 48, or 72 h). The resulting senescent cells were assessed by SA-β-gal staining, whilst cell survival and apoptosis were also analyzed by an MTT assay. Secondly, D-galactose (D-gal)-induced cell senescence model (i.e., Model 2) was established for the parallel experiments. Briefly, various concentrations of D-gal (1, 5, 10, 20, 30, 40, or 50 mg/ml) were added into the indicated cell culture media and maintained for distinct lengths of time (24, 48, or 72 h) so as to induce cellular senescence. The optimal concentrations of D-gal for senescence induction and its inducible time points were determined by both SA-β-gal and MTT assays.

### 2.5 Cell proliferation and colony formation assays

Experimental cells were allowed for growth in 96-well culture plates and treated, according to the protocol of Cell Counting Kit-8 (CCK-8) (10 μL, Solarbio), for 2 h in an 5% CO2 incubator of 37 ° C, before OD of each well were recorded at 450 nm. The resulting data were calculated to estimate cell proliferation. To assess the cell colony formation, about 500 of experimental cells were allowed for growth for 24 h in complete media in each well of 6-well culture plates, and then subjected to the required treatments for 10∼14 d. These cell culture media were changed by same fresh media on every third day. The cell colonies were fixed for 10 min by 4% of paraformaldehyde (Sangon, Shanghai, China) and then incubated for additional 20 min with 1% of the crystal violet stain reagent (Solarbio) at room temperature, followed by being rinsed with PBS twice. The staining cell colonies were photographed and counted.

### 2.6 Flow cytometric analyses of cell cycle and intracellular ROS levels

Experimental cells were pelleted by centrifugation and washed with cold PBS. The cell pellets were fixed overnight in pre-cooled 75% ethanol. The pelleted cells were further rinsed trice with PBS and re-suspended in a staining solution (Yeasen, Shanghai, China), and then allowed for incubation in the dark for at least 30 min at room temperature. These cell cycles were meticulously evaluated utilizing a flow cytometer (Beckman Coulter, Brea, CA, USA). The resulting data were analyzed by FlowJo V10 software (BD Biosciences). In some occasions, the intracellular ROS levels were measured by DCFH-DA fluorescent probes, and analyzed by flow cytometry as described previously [31].

### 2.7 Real-time quantitative PCR (qRT-PCR)

Total RNAs were isolated by using an RNA extraction kit (Promega, Madison, WI, USA) and subsequently subjected to reverse transcription to produce cDNA was performed through the cDNA Synthesis Kit (Yeasen), all according to the manufacturers’ instructions. Quantification of relative mRNA levels was assessed using the CFX Connect real-time PCR system (Bio-Rad, Hercules, CA, USA). The reaction system of qPCR Master Mix (GoTaq®, Promega) was utilized with the indicated protocol. All reactions were conducted in triplicate in a multiplate plate (96-well, Bio-Rad), and melting curve analysis was employed to determine specific amplification. Relative mRNA expression levels were normalized to those of β-actin gene controls by utilizing the 2^−ΔΔCT^ method. The sequences of all primers used for qRT-PCR are listed in Supplementary Table 1.

### 2.8 Western blot analysis

Experimental cells were washed twice with PBS and then harvested in a lysis buffer (Yeasen). The total proteins were extracted as performed at 4 °C and quantified by using another BCA kit (Yeasen). Subsequently, the protein samples were separated by SDS-polyacrylamide gel electrophoresis (PAGE) containing 8∼15% polyacrylamide, followed by being transferred onto a polyvinylidene fluoride membranes (Thermo). The protein-transferred membranes were blocked in a fast blocking buffer (Yeasen) and then subjected to immunoblotting: appropriate primary antibodies were incubated overnight at 4 °C, followed by the indicated secondary antibodies (horseradish peroxidase-conjugated; Sangon) were incubated for 2 h at room temperature. The immunoblotted protein bands were visualized with an chemiluminescent western blot detection kit (Yeasen), and their intensity was quantified by utilizing Bright Analysis software (Thermo). β-actin was employed as an internal loading control. All antibodies used here were listed in Supplementary Table 1.

### 2.9 Small interfering RNA (siRNA) transfection

Experimental cells were transfected with each of indicated siRNAs specifically targeting human Nrf2, STAG2, SMC3 and a scrambled siRNA control (Tsingke, Beijing, China, in Supplementary Table 1) in Lipofectamine™ 3000 (Thermo) according to the manufacturer’s protocol. The transfected cells were allowed for 24-h recovering growth before being subjected to indicated treatments for further experimental examinations.

### 2.10 Bioinformatics analysis of RNA sequencing

The total RNAs from *WT* and *Nrf1α*^−/−^ HeLa cell lines were isolated and purified with an RNA extraction kit (Promega), and then subjected to transcriptomic sequencing at the Beijing Genomics Institute (Shenzhen, China). Additional RNA-sequencing data from *WT* and *Nrf1α*^−/−^ HepG2 and HEK293 cells, as well as from *Nrf1α*-expressing HEK293 cells were obtained from our previous studies [30, 32]. To evaluate all reads for filtering, they were all aligned to the Refseq Human Genome GRCh38.p12 by utilizing both Bowtie2 [33] and HISAT softwares [34]. Indicated gene expression values were determined using RNA-Seq by Expectation-Maximization (RSEM) [35]. Amongst them, significantly differentially expressed genes (DEGs) were further identified by two parameters exhibiting a false discovery rate < 0.001 and | log_2_(fold changes) | >1. The DAVID tool was also employed to perform Gene Ontology annotation enrichment analysis of those selected DEGs. In addition, a protein-protein interaction (PPI) network of the DEGs was generated utilizing the STRING database. Such key nodes (hub genes) in the PPI network were identified by using the Cyotoscape plugin cytoHubba and MCODE [36, 37]. The significant hub genes were further identified through cross-comparison between datasets and also validated via qRT-PCR.

### 2.11 Luciferase reporter assay

The ARE-driven reporter and its mutants from indicated genes were constructed by subcloning relevant sequencesinto a PGL3-promoter vector. The resulting *ARE-Luc* reporter constructs or their mutants were co-transfected into HepG2 cells together with either an Nrf1-expressing construct or empty pcDNA3.1 vector control. The *pRL-TK* plasmid, coding for Renilla luciferase, served as an internal control for the transfection. After transfection for 24 h, the ARE-driven luciferase activity was determined by a dual-reporter assay kit (Promega) and then normalized to the corresponding values of Renilla luciferase.

### 2.12 Chromatin immunoprecipitation (ChIP)-sequencing analysis

The date for ChIP-sequencing were acquired from the ENCODE ChIP-Seq repository (https://www.encodeproject. org). The project identifier for the ENCODE study is ENCSR543SBE (Nrf1/Nfe2l1 in Homo sapiens HepG2 cells).

### 2.13 Statistical analysis

All relevant experimental data are displayed in the form of Mean± SD, which were obtained from at least three independent experiments. The statistical analysis of data was conducted by Student’s t-test, Mann-Whitney test or two-way ANOVA with the SPSS 22.0 software (IBM, Armonk, NY, USA). A statistically significant disparity was considered at probability (*P*) values < 0.05.

## 3 Results

### 3.1 Loss of Nrf1α enhances SA-β-gal activity in two senescence models, but inhibits proliferation of *Nrf1α*^−/−^ cells

To evaluate the role of Nrf1α in cellular senescence induced by oxidative stress, two distinct models of cell senescence were established: a simplified senescence model (i.e., Model 1) and another D-gal-induced senescence model (i.e., Model 2), as shown in Fig. S2 (A to F). These two models were assessed by integrating both outcomes of SA-β-gal staining and MTT assay, revealing that a high degree of cellular senescence is accompanied by relative low lethality (Fig. S2). As compared with *WT* cell lines, deletion of *Nrf1α* caused significant increases in the intracellular ROS levels in *Nrf1α*^−/−^ cell lines derived from HepG2 and HeLa (Fig. S3A, *Model 1*), and further incremented ROS were also stimulated by D-gal (Fig. S3A, *Model 2*). This suggests that such oxidative stress caused by *Nrf1α*^−/−^ is likely involved in induction of cell senescence, because our prior work had shown that the cellular redox homeostatic balance is disrupted by the loss of *Nrf1α,* leading to significantly elevated levels of endogenous oxidative stress [38].

In the above oxidative stress-induced senescence models, *Nrf1α*^−/−^ cells exhibited obviously enlarged and flattened senescent cell morphologies, when compared to those of *WT* cells (as illustrated in Fig. 1, A & B). Such a loss of *Nrf1α* caused a notably higher positivity in the SA-β-gal staining *Nrf1α* ^−/−^ cells than that of *WT* cells (Fig. 1, C and D). Consistent with these findings, the enzymatic activity of β-gal activity measured in both senescence models of *Nrf1α*^−/−^ cells was, *de facto*, markedly increased by comparison with that of *WT* cells (Fig. 1, E and F). Collectively, these demonstrate that the loss of Nrf1α does not only result in heightened oxidative stress in two distinct senescence models of *Nrf1α*^−/−^ cells, and also potentially exacerbate such oxidative stress-induced senescence.

**Fig. 1.**
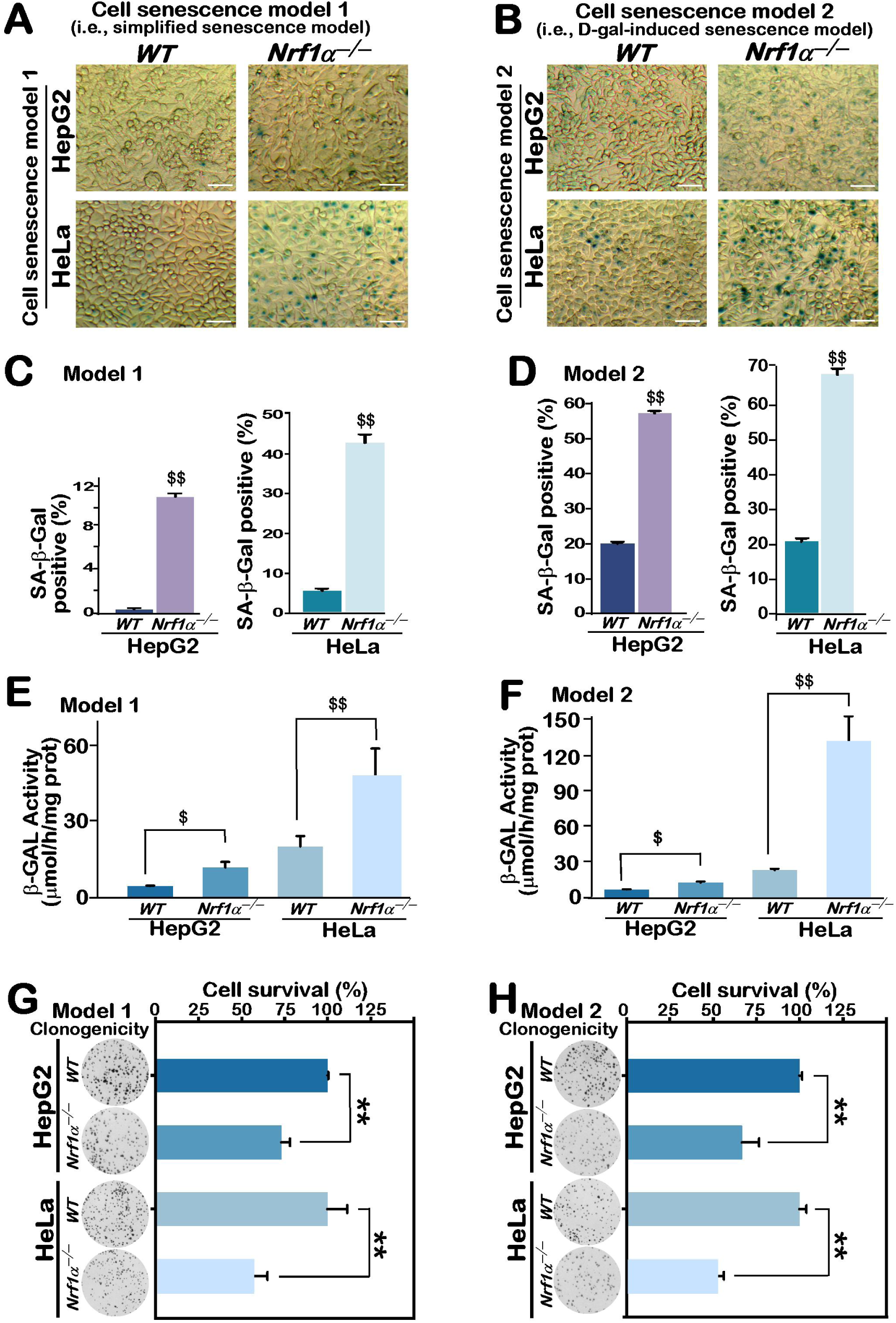
Loss of Nrf1α enhances SA-β-gal activity in two senescence models, but inhibits proliferation of *Nrf1α*^−/−^ cells. (A,B) The SA-β-gal staining of two distinct models of cellular senescence, that was established on the base of *WT* (HepG2 and HeLa) and derived *Nrf1α*^−/−^ cell lines, as described in the section of ‘Experimental procedures’. The representative images are shown herein, which were obtained from cellular senescence Model 1 (A) and Model 2 (B), respectively. Each bar = 50 μm. (C,D) The percentage of those positive SA-β-gal staining cell counts of *WT* and *Nrf1α*^−/−^ in both Model 1 (C) and Model 2 (D), as shown as Mean ± SD, together with significant increases ($$, p < 0.01), which were statistically determined from at least three independent experiments performed in triplicates (n = 3 × 3). (E,F) Quantitative analysis of the SA-β-gal activity, by measuring the generation rate of p-nitrophenol in *WT* and *Nrf1α*^−/−^ cell lines under distinct senescent conditions of Model 1 (E) and Model 2 (F). The resulting data are shown as Mean ± SD, with significant increases ($, p < 0.05; $$, p < 0.01), which were statistically determined from at least three independent experiments performed in triplicates (n = 3 × 3). (G,H) Comparative analysis of *WT* and *Nrf1α*^−/−^ cell colony formation (*left panels*) and their survival rates (*right panels*) in two distinct conditional senescence models. The latter data are shown as Mean ± SD, with significant decreases (**, p < 0.01), which were statistically determined from at least three independent experiments performed in triplicates (n = 3 × 3).

Since senescent cells characteristically exhibit impaired proliferation, it is inferable that their proliferative capacity is much likely to be negatively correlated with their senescence extents to what these cells have undergone *per se*. Thereby, we proceeded to compare distinctions in the proliferative capacity of both *WT* and *Nrf1α* ^−/−^ cellular senescence models by using their colony formation assays. The results revealed that such a lower number of colonies was caused by *Nrf1α*^−/−^ cell, when compared to those derived from *WT* cells (in *left panels* of Fig. 1, G and H). This was, also in the meantime, accompanied by a greater reduction in the survival of *Nrf1α*^−/−^ cells (in *right panels* of Fig. 1, G and H). Taken altogether, these findings indicate that deletion of Nrf1α promotes oxidative stress-induced cellular senescence phenotypes in two distinct model cell lines.

### 3.2 Deletion of Nrf1α promotes SASP of *Nrf1α*^−/−^ cells with its cycle arrest in two senescence models

Those autocrine and paracrine abilities of senescent cells are clearly known to be acquired by producing a plethora of pro-inflammatory cytokines, proteases and other relevant molecules, all of which are collectively referred to as the SASP [7, 39]. These SASP factors, thereby, play pivotal roles in the secondary senescence to drive self-senescence *via* an autocrine feedforward loop, and even induce dysfunction of neighboring cells and their growth arrest [40]. The relative expression profiling of those canonical SASP genes in two distinct cellular senescence models was examined by real-time qPCR in *WT* and *Nrf1α*^−/−^ cell lines. As shown in Fig. 2 (A–G), mRNA expression levels of *IL-1α, IL-1β, IL-6, IL-8, CXCL-3, CXCL-10* and *MMP1* were significantly up-regulated in two distinct *Nrf1α*^−/−^ cell lines compared to *WT* controls. This result indicates that deficiency of Nrf1α promotes the transcriptional expression of those core SASP factors.

**Fig. 2.**
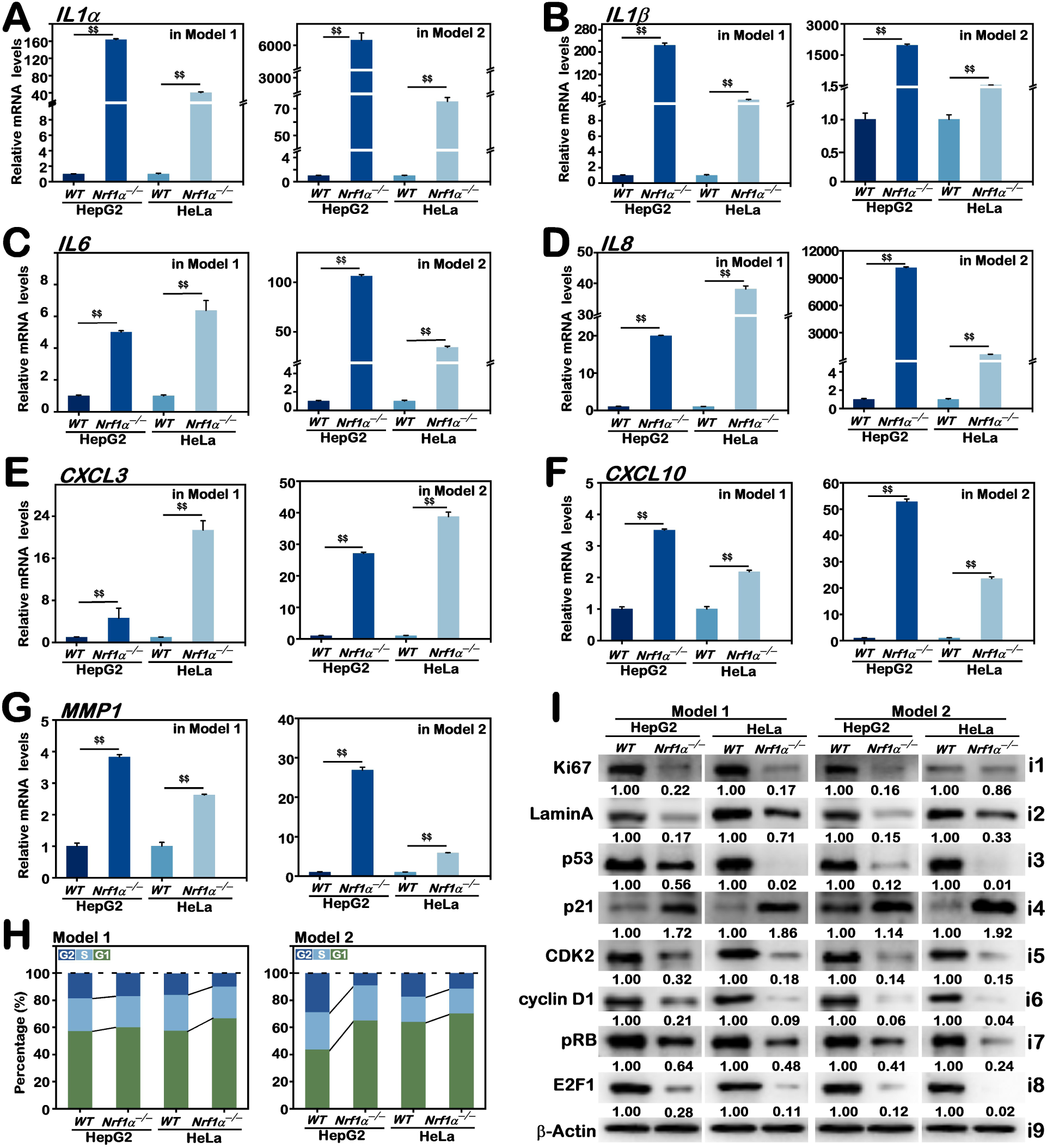
Deletion of Nrf1 promotes SASP of *Nrf1α*^−/−^ cells with its cell cycle arrest in two distinct senescence models. (A–G) The mRNA expression levels of those SASP-specific markers, *IL-1α* (A)*, IL-1β* (B)*, IL-6* (C)*, IL-8*(D)*, CXCL-3*(E)*, CXCL-10* (F), and *MMP1* (G) in *WT* and *Nrf1α*^−/−^ cell lines under distinct senescence conditions were quantified by RT-qPCR. The resulting data are shown as Mean ± SD, with significant increases ($$, p < 0.01), that were statistically determined from at least three independent experiments performed in triplicates (n = 3 × 3). (H) The protein expression abundances of Ki67, LaminA, p53, p21, CDK2, cyclin D1, pRB, and E2F1 in distinct senescent model cells of *WT* and *Nrf1α*^−/−^ were analyzed by Western blotting. The intensity of all immunoblots was quantified and normalized by that of β-actin, before being shown as fold changes (*on the bottom* of indicated blots). The data presented here were a representative of three independent experiments. (I) Changes in distinct model senescence cell cycles of both WT and Nrf1α−/− lines were determined by flow cytometry. The resulting data are shown graphically as a Mean value of three independent experiments (n = 3), of which the representative images were presented in Figure S3B.

Since a proliferation marker protein Ki67 and another nuclear membrane protein LaminA are frequently employed as two negative molecule markers of senescent cells, we hence estimated their protein expression abundances in distinct cellular senescence models of both *WT* and *Nrf1α*^−/−^ cell lines. As anticipated, the protein levels of Ki67 and LaminA were substantially down-regulated in *Nrf1α*^−/−^ cells, by contrast to those of *WT* cells (Fig. 2I, *i1* and *i2*). This result suggests a possibility that *Nrf1α*^−/−^ cells are much likely to be arrested at the G1 phase of its senescence model cell cycles. In fact, the loss of *Nrf1α*^−/−^ resulted in a marked elevation in the G1-phase cell population, whilst the proportions of both S- and G2/M-phase cells were significantly decreased, when compared to *WT* controls, in two distinct senescence cell models (Figs. 2H and S3B). This finding demonstrates deletion of Nrf1α reinforces the senescence cell cycle arrest at its G1 phase (as another crucial trait of cellular senescence). Of note, oxidative stress-induced senescence is a progressive transform from a pre-senescent stage through early senescent transition to a completely full senescence state, along as accompanied by chromatin remodeling and SASP production [7, 15]. In such progressive process, a pivotal saddle point occurs only when cells shifted from a transient state to a stable cell cycle arrest of senescence [41]. The cell cycle arrest is further believed to be facilitated by elevated DNA damage signaling caused by oxidative stress, which hinders the advancement of the cell cycle by predominant regulatory factors, e.g., retinoblastoma-associated transcriptional co-repressor(RB) at the G1/S checkpoint [42]. Next, we evaluated the protein expression levels of critical cell cycle regulatory components, such as RB, in both senescence models of *WT* and *Nrf1α*^−/−^ cell lines. Intriguingly, although p53 was substantially down-regulated or even abolished in *Nrf1α*^−/−^ cells (Fig. 2i, *i3*), its downstream regulator p21 exhibited a remarkably elevated expression in *Nrf1α*^−/−^ cells, as compared to the counterparts expressed in *WT* cells (Fig. 2I, *i4*). However, the protein levels of CDK2 and cyclin D1, which are two principal modulators of the G1/S checkpoint in the cell cycle, were significantly diminished or almost completely abolished in distinct cell lines of *Nrf1α*^−/−^(Fig. 2I, *i5* and *i6*). Such a decrease in the activation of RB kinase led to significant inhibition of RB phosphorylation (pRB) (Fig. 2I, *i7*), as accompanied by a substantial attenuation or even abolishment of the transcription factor E2F1 expression levels in *Nrf1α*^−/−^ cells (Fig. 2I, *i8*), ultimately resulting in an intensified cell cycle arrest of *Nrf1α*^−/−^ at its G1/S checkpoint in two distinct senescence models. These findings suggest that the loss of Nrf1α leads to the evident dysregulation of key cell cycle regulatory factors responsible for its G1/S checkpoint, consequently exacerbating senescence cell cycle arrest. Overall, these collective data regarding alterations attributable to the cellular senescence process of *Nrf1α*^−/−^ cells in two distinct senescence models induced by oxidative stress, together with those key senescence markers, demonstrate that oxidative stress induced-senescence is exacerbated by Nrf1α deficiency.

### 3.3 Cell senescence is exacerbated by *Nrf1α*^−/−^ disruption of its STAG2/SMC3-dependent chromosomal homeostasis

Based on the above-described experimental evidence, the indispensable redox-determining factor Nrf1 is postulated to exert its significant anti-senescence function in maintaining cellular homeostatic equilibria, as proposed previously [20, 22, 38]. Those differentially expression genes (DEGs) regulated by Nrf1 critically responsible for maintaining cellular homeostasis and preventing cellular senescence are herein identified through transcriptome sequencing analysis. After normalizing the gene expression datasets, screening and conducting functional enrichment analysis of DEGs, we further identified 18 candidate crucial genes through a PPI network and module analysis (Fig. 3A). Subsequently, we conducted further core analysis and, followed by real-time qPCR validation (Fig. S4A), narrowed down our focus on two key targets, namely, STAG2 and SMC3 (Fig. S4B), as two components of the cohesin complex, which are primarily associated with the chromosomal homeostasis [43], because they participate in preserving the chromosomal number and structure integrity, ensuring chromosomal stability and thereby play crucial important roles in the gene expression and DNA repair. It is well-established that STAG2 and SMC3 are essential for cell viability, but conversely, substantially down-regulated expression of STAG2 and SMC3 in *Nrf1α*^−/−^ cells as compared to *WT* controls (Fig. 3B) are much likely to result in a marked decrease in its viability.

**Fig. 3.**
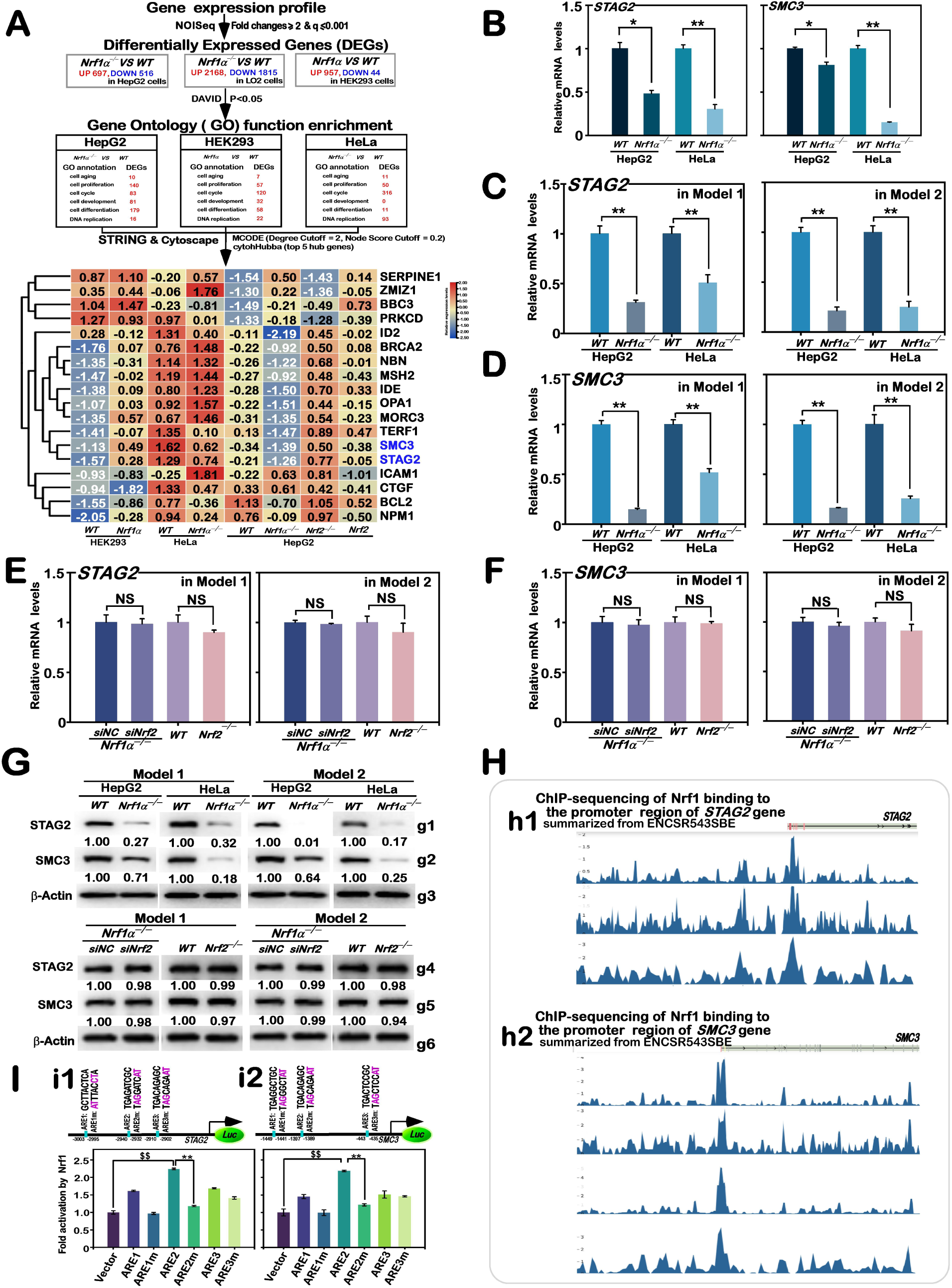
STAG2 and SMC3 are identified as two direct target genes of Nrf1 but not Nrf2 in the anti-senescent responses. (A) A flow of bioinformatics analysis of those transcriptome sequencing data from distinct cell lines (*WT*, Nrf1 α -expressing, Nrf2-activating, *Nrf1α*^−/−^ or *Nrf2*^−/−^), as shown by key differentially expressed genes (DEGs) regulated by Nrf1 that are selected critically for maintaining cellular homeostasis in putative anti-senescent response. A group of selected candidate genes were indicated as distinct expression levels in the heatmap, in which *SMC3* and *STAG2* were highlighted (*in blue*). (B) Real-time qPCR analysis of *STAG2* and *SMC3* at their mRNA levels expressed in *WT* and *Nrf1α*^−/−^ cell lines, that had been cultured under normal conditions. The resulting data are shown as Mean ± SD, with significant decreases (*, p < 0.05; **, p < 0.01), which were statistically determined from at least three independent experiments performed in triplicates (n = 3 × 3). (C,D) Under two distinct senescence model conditions, the mRNA expression levels of *STAG2* (C) and *SMC3*(D) in *WT* and *Nrf1α*^−/−^ cell lines were determined by real-time qPCR. The resulting data are shown as Mean ± SD, with significant decreases (**, p < 0.01), which were statistically determined from at least three independent experiments performed in triplicates (n = 3 × 3). (E,F) Real-time qPCR analysis of *STAG2* (E) and *SMC3* (F)at their mRNA levels expressed in distinct HepG2 cell lines represented by *WT*, *Nrf2*^−/−^, *Nrf1α*^−/−^+*siNC*, and *Nrf1α*^−/−^+*siNrf2*, all of which were placed in two distinct cell senescence conditions. The resulting data are shown as Mean ± SD, with no statistical differences (NS), which were determined from at least three independent experiments performed in triplicates (n = 3 × 3). (G) Western blotting analysis of the protein levels of STAG2 and SMC3 in the indicated cell lines of *WT* and *Nrf1α*^−/−^ (derived from HepG2 and HeLa cells), along with another group of *WT*, *Nrf2*^−/−^, *Nrf1α*^−/−^+*siNC*, and *Nrf1α*^−/−^+*siNrf2* (derived from HepG2 cells), all of which were cultured in two distinct cell senescence conditions. The intensity of all immunoblots was quantified and normalized by that of β-actin, before being shown as fold changes (*on the bottom* of indicated blots). The data presented herein were a representative of three independent experiments. (H) The ChIP-seq data of Nrf1-binding to the gene promoters of STAG2 (h1) and SMC3 (h2) in WT HepG2 cells, which were obtained from the Encode platform with relevant information. (I) Two schematic representations of distinct luciferase reporters driven ARE sequences from both *STAG2* (i1) and *SMC3* (i2) gene promoter regions and their corresponding mutants (*Upper panels*). Subsequently, HepG2 cells were co-transfected with the indicated ARE-driven reporters or mutants, plus another internal control pRL-TK, along with an Nrf1 expression construct or an empty pcDNA3 vector. The ARE-driven Luciferase activity regulated by Nrf1 was calculated and graphically shown as Mean ± SD (*lower panels*), relative to their background activity measured by empty pcDNA3 control. Significant increases ($$, p < 0.01) and significant decreases (**, p < 0.01) were statistically determined from at least three independent experiments performed in triplicates (n = 3 × 3).

To further clarify whether STAG2 and SMC3 are involved in the cellular senescence enhanced by *Nrf1α*^−/−^-leading oxidative stress, we evaluated their mRNA expression in both *WT* and *Nrf1α*^−/−^ cell lines in two distinct senescence cell models. As shown in Fig. 3(C and D), a dramatically down-regulation profiling of STAG2 and SMC3 at their mRNA levels was observed in *Nrf1α*^−/−^ cell lines when compared to their WT controls. Similarly, abundances of both STAG2 and SMC3 proteins were also substantially down-regulated in *Nrf1α*^−/−^ cells (Fig. 3G, *g1* and *g2*). Such mRNA and protein expression levels of both STAG2 and SMC3 in *Nrf1α*^−/−^ cells were almost unaffected by silencing of *siNrf2* in (Fig. 3E, 3F, and 3G, *g4* and *g5*), and no significant changes in their basal expression levels were observed in *Nrf2*^−/−^ cells. Therefore, these collective results demonstrate that the loss of Nrf1α, but not of Nrf2, predominantly down-regulates the expression of STAG2 and SMC3 in two distinct cell senescence models. Further examinations revealed that significant increases in the transactivation activity of those indicated *ARE*-driven luciferase reporters from both *STAG2* and *SMC3* genes (particularly STAG2/SMC3-ARE2-luc) was directly regulated by Nrf1 in *WT* HepG2 cells (Fig. 3I). In addition, the DNA-binding activity of Nrf1 in the *STAG2* and *SMC3* promoter regions was further validated in HepG2 cells by relevant ChIP-sequencing date (Fig. 3H, obtained from the ENCODE repository (https://www.encodeproject.org)).

Next, to confirm putative roles of STAG2 and SMC3 in the cellular senescence, their specific siRNAs were transfected into *WT* HepG2 cells, before the cell senescence markers were assessed (Fig. 4). As anticipated, significant increases in the senescence-activated β-gal activity were caused by silencing of *siSTAG2* or *siSMC3* in two distinct cell models, when compared to its *siNC* controls (Fig. 4A). Furtherly, all examined SASP factors, *IL-1α, IL-1β, IL-6, IL-8, CXCL-3, CXCL-10*, and *MMP1* at their mRNA expression levels in two distinct models of cellular senescence (Fig. 4, B to H) were significantly up-regulated by *siSTAG2* or *siSMC3* (Fig. 4I, *i1*). Additional two key senescence biomarker Ki67 and LaminA also exhibited sharp decreases in their protein abundances upon silencing of STAG2 or SMC3 (Fig. 4I, *i2* and *i3*). Besides, the cell cycle regulatory factors p53 and p21 showed remarkable enhancements (Fig. 4I, *i4* and *i5*), whereas CDK2, cyclin D1, pRB, and E2F1 were dramatically downregulated in either STAG2 or SMC3-silenced cells, as compared with their controls in two distinct senescence cell models (Fig. 4I, *i6* to *i9*). Collectively, these results demonstrate that the silencing of STAG2 or SMC3 leads to an acceleration of cellular senescence. Overall, the deletion of Nrf1 results in exacerbated cell senescence induced by oxidative stress primarily through disrupting the homeostatic expression of STAG2 or SMC3, hence hindering the formation of chromosome cohesion and also causing chromosomal instability (Fig. 4J). Such a disturbance of the cellular chromosomal homeostasis caused by *Nrf1α*^−/−^-leading dysfunctional expression of its downstream targets STAG2 or SMC3 leads ultimately to accelerating the cellular senescence induced by oxidative stress

**Fig. 4.**
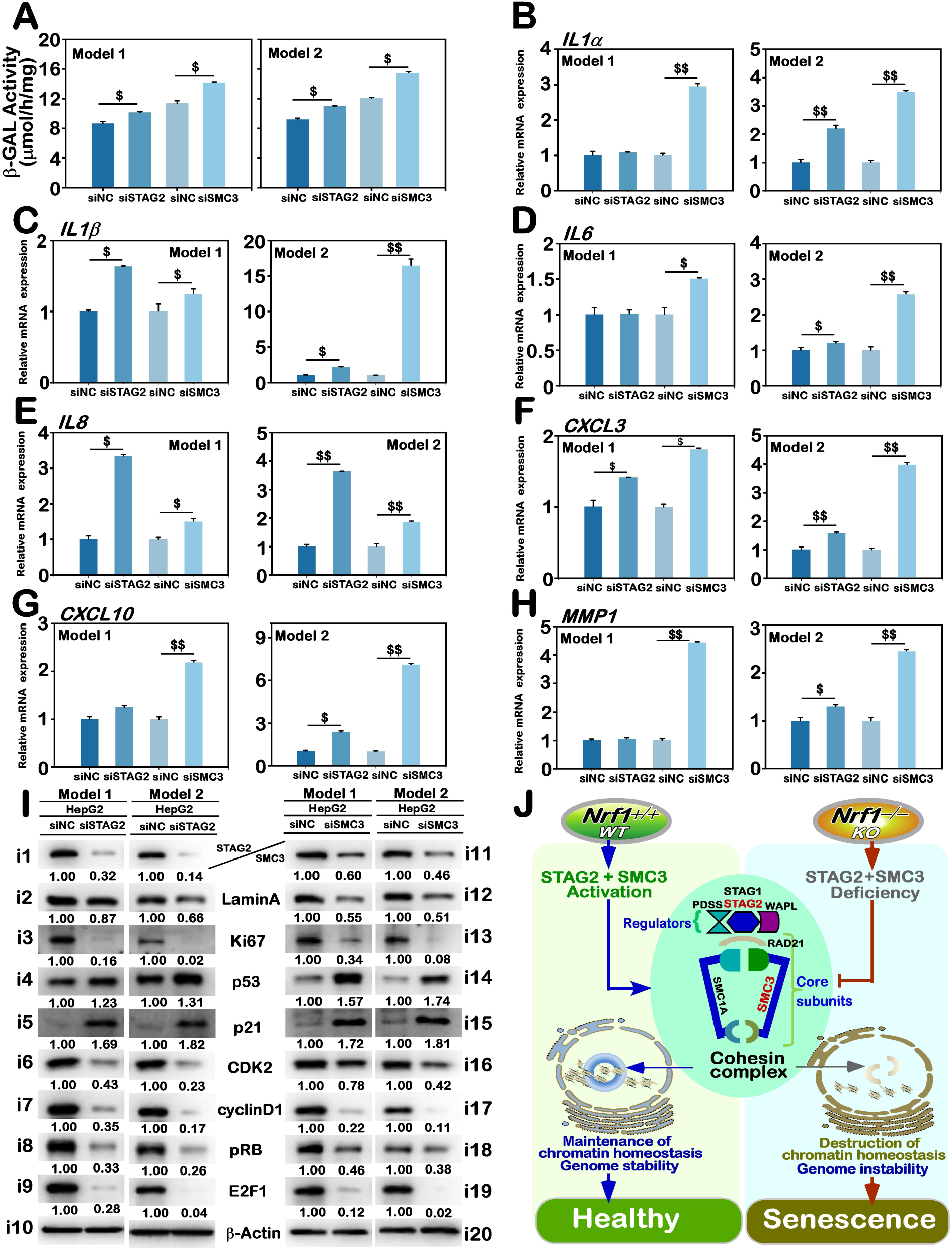
Cellular senescence enhanced by silencing of *STAG2* and *SMC3* required for chromosomal homeostasis. (A) The SA-β-gal activity was determined by its enzymatic quantitative assay in HepG2 cells where *STAG2* or *SMC3* had been knocked down respectively by *siSTAG2* or *siSMC3* in two distinct senescence cell models. The resulting data are shown as Mean ± SD, with significant increase ($, p < 0.05), which were statistically determined from at least three independent experiments performed in triplicates (n = 3 × 3) (B–H) The mRNA expression levels of SASP-specific markers, *IL-1α* (B)*, IL-1β* (C)*, IL-6* (D)*, IL-8* (E)*, CXCL-3* (F)*, CXCL-10* (G), and *MMP1* (H) in *siSTAG2-* or *siSMC3*-silenced cell lines under distinct model senescence conditions were determined by real-time qPCR. The resulting data are shown as Mean ± SD, with significant increases ($, p < 0.05; $$, p < 0.01) or no statistical differences (NS), which were statistically determined from at least three independent experiments performed in triplicates (n = 3 × 3). (I) The protein expression levels of Ki67, LaminA, p53, p21, CDK2, cyclin D1, pRB, and E2F1 in the above-described *siSTAG2-* or *siSMC3*-silenced cell lines were analyzed by Western blotting with distinct antibodies. The intensity of all immunoblots was quantified and normalized by that of β-actin, before being shown as fold changes (*on the bottom* of indicated blots). The data presented herein were a representative of three independent experiments. (J) A diagram depicting the putative mechanism by which Nrf1-targets STAG2 and SMC3 in the cohesion complex are required for maintaining cellular chromosomal homeostasis in the anti-senescence response.

### 3.4 Cellular senescence is exacerbated by *Nrf1α*^−/−^-leading dysfunction of autophagy in two distinct cell models

There exists a growing body of consensus suggesting that the development of cellular senescence is associated with dysregulation of autophagy, particularly under oxidative stress [44, 45]. As a crucial elemental factor governing cellular homeostasis, cell autophagy is also a continuous process, encompassing the creation of autophagosomes, its subsequent merging with lysosomes and ensuing degradation, and its unique functioning is enabled through sequential involvement of multiple functional complexes consisting of those core autophagy-related (ATG) proteins. It is widely agreed upon that all those ATG core molecules are indispensable for autophagy, because the normal expression of each autophagy core molecule is imperative for the effective operation of autophagy [46]. Herein, we have investigated potential impacts of *Nrf1α* ^−/−^ deficiency on autophagic capability in oxidative stress-induced senescence models, by real-time qPCR analysis of the mRNA expression levels of those core autophagy markers. The results revealed that *ULK1*, *Beclin1, ATG4A, ATG7, ATG9A, ATG10,* and *ATG12* were markedly down-regulated in two distinct models of *Nrf1α*^−/−^ cell lines, in comparison to their counterpart controls (Fig. 5, A to G). Further investigation of the protein expression of core autophagy markers by Western blotting also unraveled remarkable reductions in the basal abundances of ULK1, ATG13, Beclin1, ATG9A, ATG5, ATG10, and ATG7 in *Nrf1α*^−/−^ cells (Fig. 5H, *h1* to *h7*). These findings indicate that the loss of Nrf1 significantly restricts the expression of autophagy core markers and may thus result in impaired autophagy of indicated cellular senescence models. This is supported by further experimental evidence showing that abundances of microtubule-associated protein 1 light chain 3 (LC3)-II, representing the quantity of autophagosomes formed, and the selective autophagic receptor p62 (also called sequestosome 1, SQSTM1, which directly binds to LC3) were markedly enhanced in *Nrf1α* ^−/−^ cell lines, but the LC3-I was down-expressed, when compared to their *WT controls* (Fig. 5H, *h8* and *h9*). This suggests that the lack of Nrf1 cannot only impedes the creation of autophagic vesicles and also interferes with their fusion with the lysosome and ensuing degradation, leading to the buildup of p62 and LC3-II within the cell (Fig. 5I). Based on these, it is inferable that the deletion of Nrf1 is much likely to exacerbate the inhibition of autophagy occurring in distinct cellular senescence models induced by oxidative stress.

**Fig. 5.**
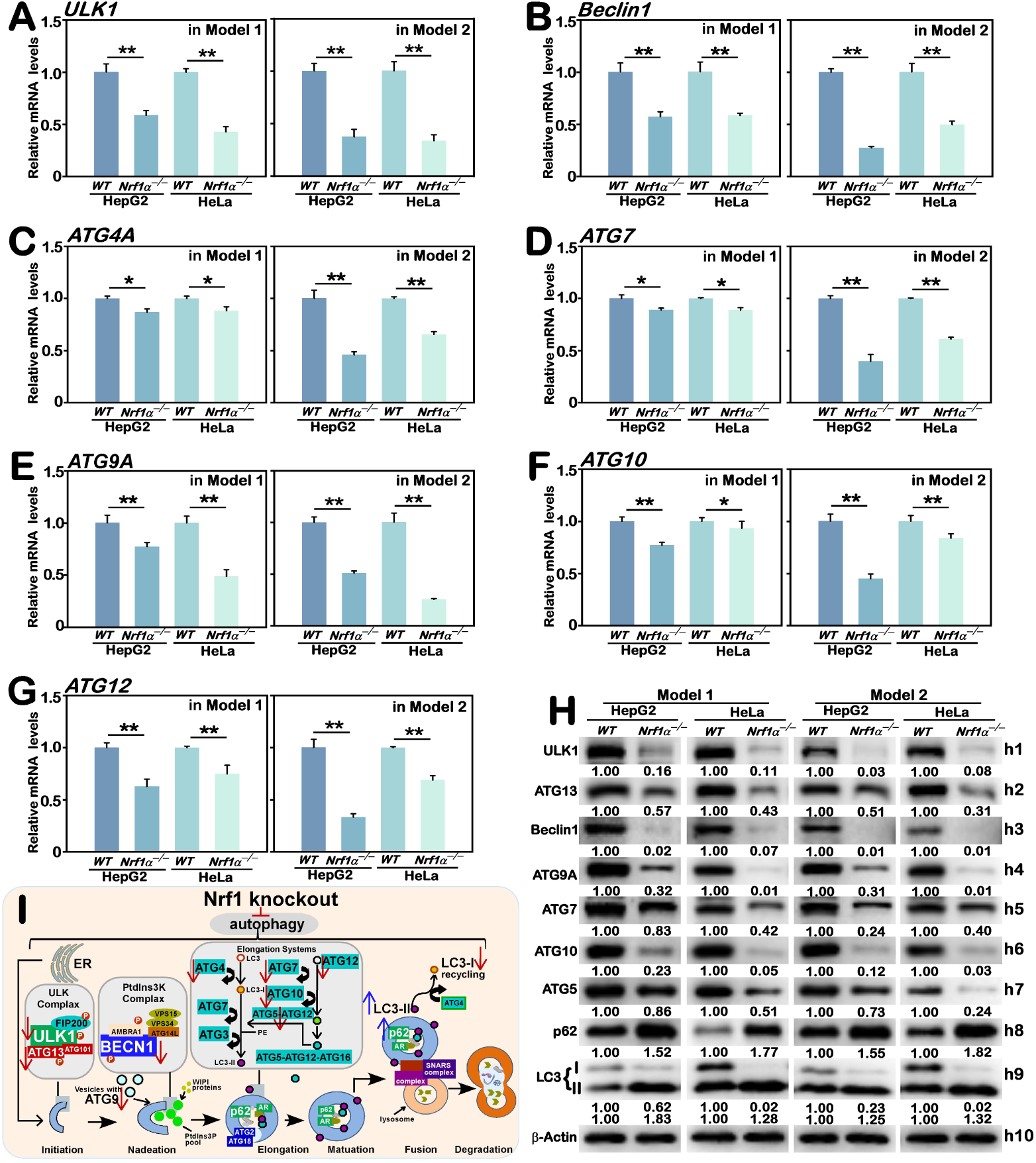
Deletion of Nrf1 leads to enhanced dysfunction of autophagy in *Nrf1α*^−/−^ cellular senescence models. (A–G) The mRNA expression levels of autophagy-related biomarkers *ULK1* (A)*, Beclin1* (B)*, ATG4A* (C)*, ATG7* (D)*, ATG9A* (E)*, ATG10* (F), and *ATG12* (G) in *WT* and *Nrf1α*^−/−^ cell lines under distinct model senescence conditions were determined by real-time qPCR. The resulting data are shown as Mean ± SD, with significant decreases (*, p < 0.05; **, p < 0.01), which were statistically determined from at least three independent experiments performed in triplicates (n = 3 × 3). (H) The protein expression abundances of ULK1, Beclin1, ATG5, ATG7, ATG9A, ATG10, ATG13, p62, and LC3 in in *WT* and *Nrf1α*^−/−^ cell lines under distinct model senescence conditions were determined by Western blotting with distinct antibodies. The intensity of all immunoblots was quantified and then normalized by that of β-actin, before being shown as fold changes (*on the bottom* of indicated blots). The data presented herein were a representative of three independent experiments. (I) A schematic illustrating the putative mechanism whereby the loss of Nrf1 leads to dysfunctional autophagy in cellular senescence.

Given that cell autophagy is a recurring and dynamic process, next we examined the autophagic flux, by definition of putative changes in the amount of autophagic activity and provides a more accurate assessment of autophagic activity. Therefore, the mTOR inhibitor rapamycin (Rapa, Fig. S5) and another autophagy inhibitor chloroquine (CQ, Fig. S6) were employed to evaluate the autophagic flux of *WT* and *Nrf1α*^−/−^ cell lines within two distinct senescence models. Following treatment with Rapa, the autophagy flux displayed a characteristic pattern of increased accumulation of core molecules as a result of Rapa-stimulated expression levels (Fig. S5). Similarly, the autophagy flux following treatment with CQ also exhibited another characteristically increased pattern of the core molecule accumulation as a result of the CQ’s inhibition of their degradation (Fig. S6). The above notions were evidenced by experiments showing that significant decreases in the mRNA expression levels of *ULK1, Beclin1, ATG4A, ATG7, ATG9A, ATG10,* and *ATG12* and the protein abundances of ULK1, Beclin1, ATG5, ATG7, ATG9A, ATG10, and ATG13 were accompanied by marked enhancements of LC3-II and p62 accumulation in *Nrf1α*^−/−^ cells, by striking comparison with their counterparts measured from *WT* cells, after both model cell lines had been treated with Rapa (Fig. S5, B to I) or CQ (Fig. S6, B to I). Collectively, these indicate that deletion of Nrf1α is much likely to impede the autophagy flux, and even autophagy progression in distinct cell senescence models. It is, thereby, inferable that the loss of Nrf1 also likely caused dysfunction of cellular autophagy to be worsened in two distinct senescence models of *Nrf1α*^−/−^ cells, potentially intensifying disruption of cellular homeostasis. Conversely, the disturbance of autophagy in *Nrf1α*^−/−^ cells is also hypothesized to account for a potential mechanism contributable to its accelerated senescence induced by oxidative stress due to Nrf1α deficiency, albeit this needs to be further experimentally evidenced.

### 3.5 Accumulation of Nrf2 in *Nrf1α*^−/−^-cells only exerts a limited compensatory anti-senescence effect for loss of Nrf1α

Nrf1 and Nrf2 jointly participate in numerous physiological responses to oxidative stress, and also diversity of their effects and relevant mechanistic cross-talks in the regulatory processes are also recognized [38]. In fact, our prior studies with transcriptomic data unraveled that the absence of Nrf1 has a minimal effect on the transcriptional synthesis of Nrf2, but the abnormal buildup of Nrf2 in *Nrf1α*^−/−^ cells is attributed to the compromised degradation of this CNC-bZIP protein by the 26S proteasome [30]. In similar normal conditions, basal mRNA expression of *p62* was down-regulated to different extents in *Nrf1α*^−/−^ or *Nrf2*^−/−^ cell lines, but significantly up-regulated by the constitutive activation of Nrf2 with a deletion mutant of its N-terminal Keap1-binding domain (i.e., *caNrf2^ΔN^*) (Fig. 6A), implying the transcriptional expression of p62 monitored primarily by Nrf2, albeit also regulated by Nrf1. By contrast, mRNA levels of *Keap1* was almost unaffected by *Nrf1α*^−/−^, *Nrf2*^−/−^ or *caNrf2^ΔN^*.

**Fig. 6.**
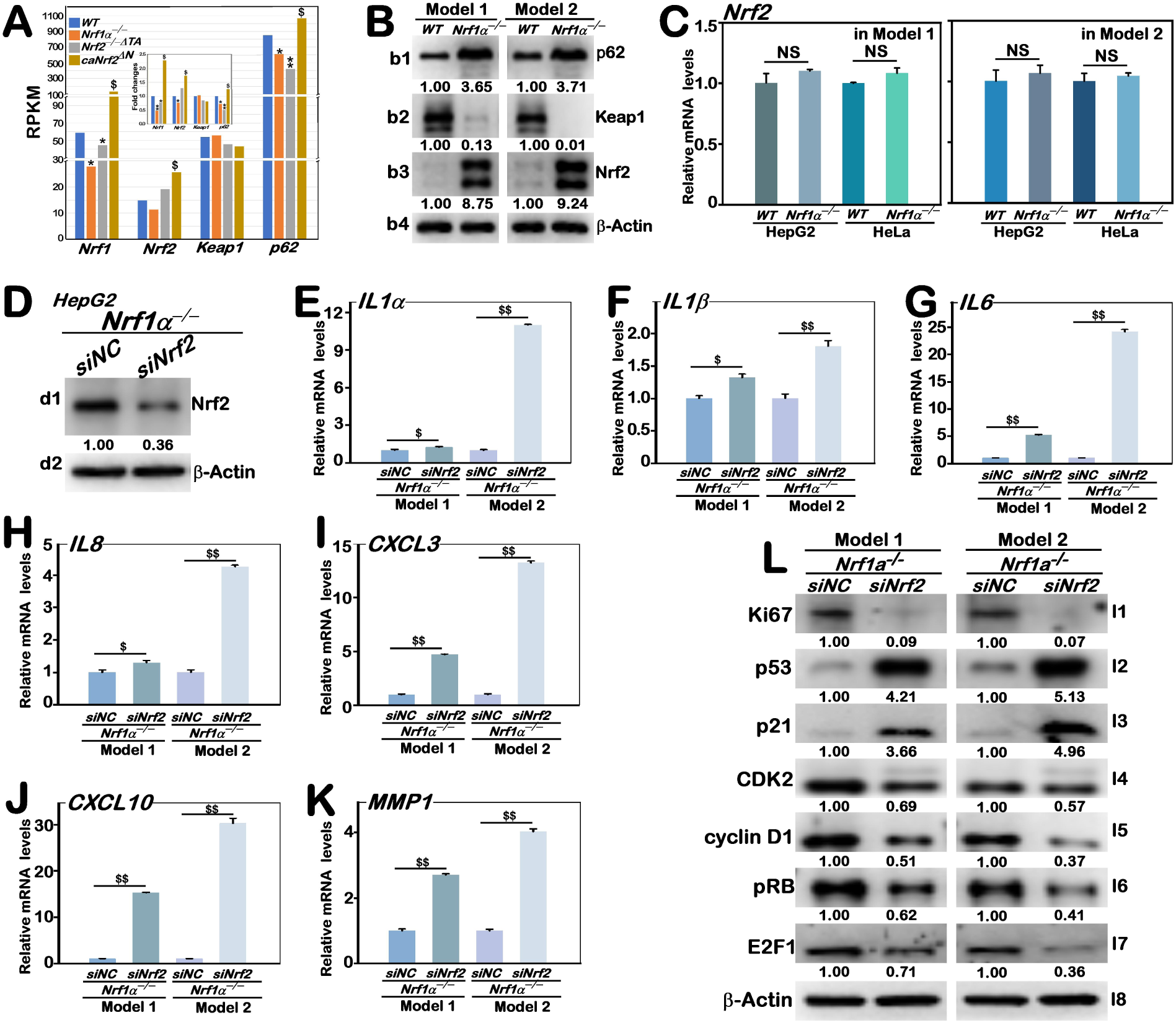
Accumulated Nrf2 in *Nrf1α*^−/−^ cells exerts a limited compensatory anti-senescence effect for such loss of Nrf1. (A) The mean RPKM mRNA values of Nrf1, Nrf2, Keap1, and p62 and their fold changes (internal panel) were calculated from transcriptome sequencing of indicated HepG2 cell lines *WT, Nrf1α*^−/−^, *Nrf2^−/−ΔTA^*, and *caNrf2^ΔN^*. The data are graphically shown, with significant increases ($, p < 0.05) and significant decreases (*, p < 0.05; **, p < 0.01), which were statistically determined from at least three independent experiments. (B) The protein expression levels of p62, Keap1, and Nrf2 in *WT* and *Nrf1α*^−/−^ cell lines under distinct model senescence conditions were determined by Western blotting with distinct antibodies. The intensity of all immunoblots was quantified and then normalized by that of β-actin, before being shown as fold changes (on the bottom of indicated blots). The data presented herein were a representative of three independent experiments. (C) The mRNA levels of Nrf2 expressed in *WT* and *Nrf1α*^−/−^ cell lines under distinct model senescence conditions were determined by real-time qPCR. The results are shown as Mean ± SD, with no significant differences (NS), which were statistically determined from at least three independent experiments performed in triplicates (n = 3 × 3). (D) Changes in the Nrf2 protein abundances in *Nrf1α*^−/−^ cells that had been silenced by siNrf2 or siNC were assessed by Western blotting. The intensity of all immunoblots was quantified and then normalized by that of β-actin, before being shown as fold changes (on the bottom of indicated blots). The data presented herein were a representative of three independent experiments. (E–K) Real-time qPCR analysis of changes in the mRNA expression of SASP-specific markers, *IL-1α* (E)*, IL-1β* (F)*, IL-6* (G)*, IL-8* (H)*, CXCL-3*(I)*, CXCL-10*(J), and *MMP1* (K), in *Nrf1α*^−/−^ cells that had been silenced by siNrf2 or siNC under distinct model senescence conditions. The resulting data are shown as Mean ± SD, with significant increases ($, p < 0.05; $$, p < 0.01), which were statistically determined from at least three independent experiments performed in triplicates (n = 3 × 3). (L) The above-described cell lines were subjected to Western blotting with distinct antibodies against Ki67, p53, p21, CDK2, cyclin D1, pRB, and E2F1. The intensity of all immunoblots was quantified and then normalized by that of β -actin, before being shown as fold changes (on the bottom of indicated blots). The data presented herein were a representative of three independent experiments.

Next, further examinations of two distinct cell senescence models revealed that the protein accumulation of p62 resulting from autophagy dysfunction in *Nrf1α*^−/−^ -leading senescent cells was accompanied by substantial diminishment or even abolishment of Keap1 protein (i.e., a negative regulator of Nrf2), as compared to *WT* controls (Fig. 6B, *cf. b1 with b2*). As a result, this is much likely to leads to the activation of the Keap1-Nrf2-p62 axis, possibly resulting in the aberrant accumulation of Nrf2 and p62 proteins (that may lead to further decreased expression of Keap1) in *Nrf1α*^−/−^ cells (Fig. 6B). However, it is important to note that no obvious changes in the mRNA expression levels of Nrf2 were determined by real-time qPCR analysis of *Nrf1α*^−/−^ cell senescence models, by comparison to *WT* cell lines (Fig. 6C). Collectively, these data indicate that the *Nrf1α*^−/−^-leading accumulation of p62 (but with no increases in its mRNA levels) cannot result from its transcriptional expression monitored by hyperactive Nrf2, that was aberrantly accumulated in severe endogenous oxidative stress-induced cellular senescence models.

The impacts of such aberrantly-accumulated Nrf2 on *Nrf1α*^−/−^ cell senescence models were interestingly investigated, because of the significant inter-regulation between both CNC-bZIP factors in fulfilling their essential physio-pathological functions at various levels [30, 38]. After the hyperactive Nrf2 abundance in *Nrf1α*^−/−^ cells was knocked down by silencing of *siNrf2* (Fig. 6D), the mRNA expression levels of all examined SASP factors *IL-1α, IL-1β, IL-6, IL-8, CXCL-3, CXCL-10*, and *MMP1* in *Nrf1α*^−/−^ cell lines were, however, significantly enhanced by *siNrf2*, when compared to *siNC* controls, in two distinct models of cellular senescence (Fig. 6, E to K). Similarly, the protein expression of Ki67 was substantially decreased by *siNrf2*, relative to its controls (Fig. 6L, *l1*). Rather, such silencing of *Nrf2* led to significantly enhanced expression levels of both p53 and p21 proteins (Fig. 6L, *l2* and *l3*), as accompanied by substantial decreases in the protein expression of CDK2, cyclin D1, pRB, and E2F1 (Fig. 6L, *l4 to l7)*. Collectively, these indicate that the *Nrf1α*^−/−^-leading cellular senescence are execrated after being treated with *siNrf2* in the extent senescence conditions, which exhibited typical features of senescent cells, such as upregulated expression of SASP, degradation of the proliferation marker Ki67, and inactivation of the p21-RB-E2F1 axis. Hence, it is inferable that the abnormal buildup of Nrf2 in *Nrf1α*^−/−^ cells may also confer a certain extent of the cellular resistance to senescence induced by oxidative stress.

Further assessment of *Nrf2*^−/−^ impact on two distinct cellular senescence models revealed significant increases in the SA-β-gal positive staining (Fig. 7A) and its quantitative enzymatic activity (Fig. 7B), when compared to their *WT* controls. This result was also accompanied by elevated mRNA expression levels of *IL-1α, IL-1β, IL-6, IL-8, CXCL-3, CXCL-10,* and *MMP1* in *Nrf2*^−/−^ cells during distinct senescence conditions (Fig. 7, C to I), indicating that the loss of Nrf2 also leads to promoted secretion of key SASP factors. In addition, a fewer of *Nrf2*^−/−^ cell colonies were formed as compared to those of *WT* cells (Fig. 7J, *left panels*), concomitantly with a decrease in its cell survival capacity (Fig. 7J, *right panels*). Flow cytometric analysis of *Nrf2*^−/−^ cell cycles during distinct senescence conditions unraveled a marked increase in its G1-phase cell population, as accompanied by another marked decrease in its S-phase cell population, upon comparison to *WT* cell controls (Fig. 7, K–L), implying that such senescent cell cycle arrest at G0/G1 phase is also likely caused by the loss of Nrf2. Furthermore, *Nrf2*^−/−^ cells gave rise to pronounced decreases in the expression levels of Ki67 and LaminA compared to *WT* cells (Fig. 7M, *mi* and *m2*), but increased expression levels of p53 and p21 (*m3, m4*). In accordance with this, markedly diminished expression levels of CDK2, cyclin D1, pRB, and E2F1 were also observed in *Nrf2*^−/−^ cells (Fig. 7M, *m5 to m8*). Collectively, these results indicate that Nrf2 deficiency can also accelerate cellular senescence induced by oxidative stress. Hence, it is inferable that Nrf2 appears, similar to Nrf1, to have a crucial anti-senescence capability to safeguard the cell survival against oxidative stress-induced senescence, but this capability is too limited to effectively compensate for the absence of Nrf1 α, even though the hyperactive Nrf2 was aberrantly accumulated in *Nrf1α*^−/−^ cells. This fact demonstrates that the limited contribution of Nrf2 to the resistance against senescence induced by oxidative stress is just supplemental, but not so indispensable as that of Nrf1. As such being the case that the elevated Nrf2 cannot fully compensate for the deficiency of Nrf1α, it is also likely to give rise to a limited slackening, as opposed to a complete halt, in accelerating oxidative stress-induced senescence caused by the deletion of Nrf1. Overall, Nrf1 possesses a unique anti-senescence characteristic, that is distinctive from that of Nrf2, as evinced differentially in two distinct models of cell senescence, particularly induced by oxidative stress.

**Fig. 7.**
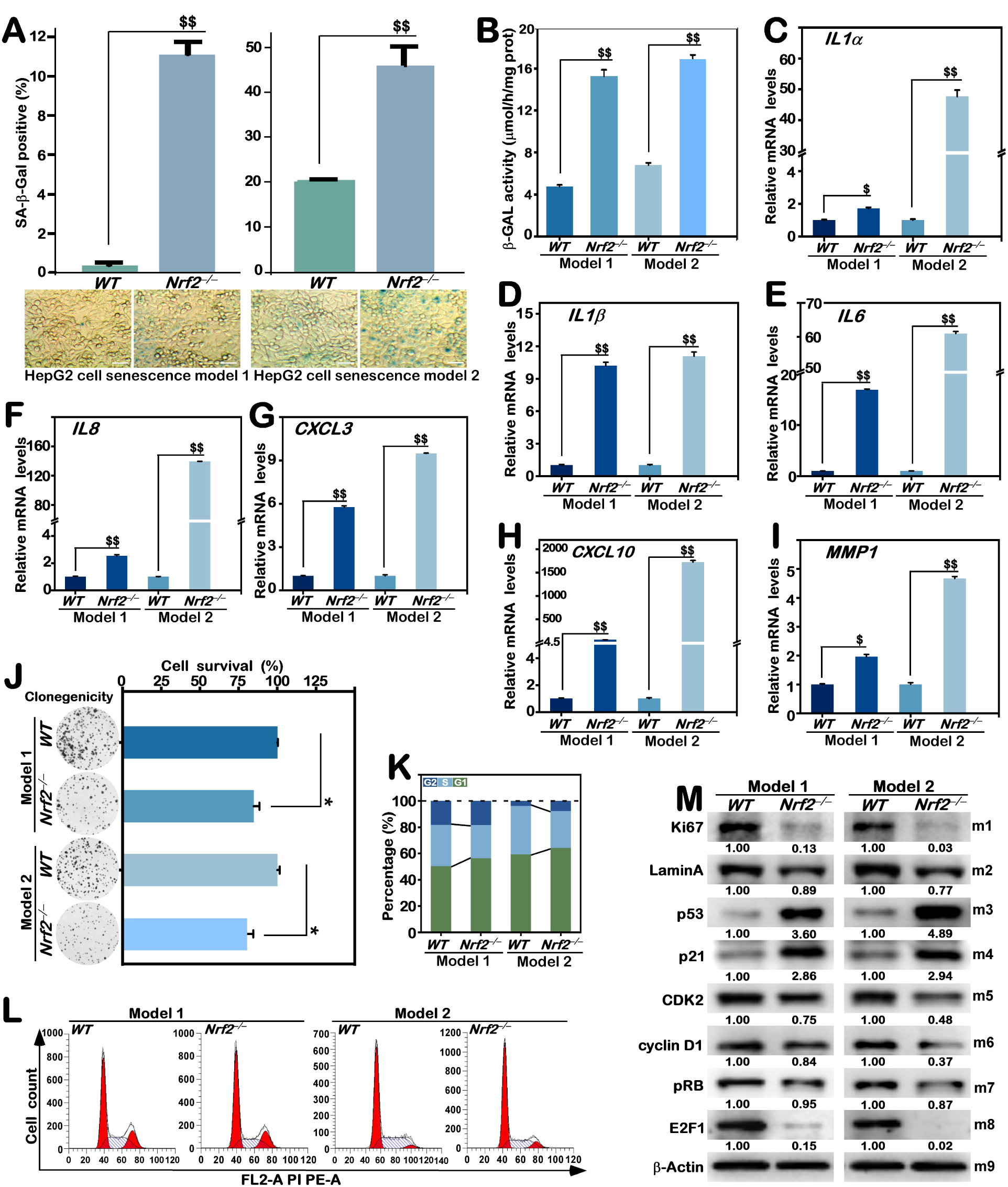
Loss of Nrf2 causes an enhancement of cellular senescence in two distinct models. (A) The SA-β-gal staining of *WT* and *Nrf2*^−/−^ cell lines in two distinct senescence models (*lower panels,* each bar= 50 μm), along with quantitative analysis of this enzymatic positive staining cells as Mean ± SD (*upper panels*), with significant increases ($$, p < 0.01), which were statistically determined from at least three independent experiments performed in triplicates (n = 3 × 3). (B) The SA-β-gal activity in *WT* and *Nrf2*^−/−^ cell lines under distinct model senescence conditions was determined by the yield of p-nitrophenol. The results are shown as Mean ± SD, with significant increases ($$, p < 0.01), which were statistically determined from at least three independent experiments performed in triplicates (n = 3 × 3). (C–I) The mRNA expression levels of SASP-specific markers *IL-1α* (C)*, IL-1β* (D)*, IL-6* (E)*, IL-8* (F)*, CXCL-3* (G)*, CXCL-10* (H)*, and MMP1* (I), in *WT* and *Nrf2*^−/−^ cell lines under distinct model senescence conditions was determined by real-time qPCR. The resulting data are shown as Mean ± SD, with significant increases ($, p < 0.05; $$, p < 0.01), which were statistically determined from at least three independent experiments performed in triplicates (n = 3 × 3). (J) Colony formation (*left panels*) of *WT* and *Nrf2*^−/−^ cell lines, along with their cell survival rate (*right panels*) were evaluated under distinct model senescence conditions. The results are shown as Mean ± SD, with significant decreases (*, p < 0.05), which were statistically determined from at least three independent experiments performed in triplicates (n = 3 × 3). (K–L) The above-described cell lines were subjected to flow cytometry analysis of these cell cycles of *WT* and *Nrf2*^−/−^. The results are shown by a percentage distribution of cell cycle phases (K) and their original flow cytometry histograms (L). (M) Changes in the protein levels of Ki67, LaminA, p53, p21, CDK2, cyclin D1, pRB, and E2F1 in *WT* and *Nrf2*^−/−^ under distinct model senescence conditions was determined by Western blotting. The intensity of all immunoblots was quantified and then normalized by that of β-actin, before being shown as fold changes (*on the bottom* of indicated blots). The data presented herein were a representative of three independent experiments.

## 4 Discussion

In the present study, we found that the loss of Nrf1α resulted in elevated senescence characteristics in *Nrf1α*^−/−^ cells under oxidative stress-induced senescence conditions, as evinced by general senescence traits and well-recognized core senescence indicators, such as heightened SA-β-gal activity, intensified cell cycle arrest and SASP. Conversely, it is inferable that such an anti-senescent role of Nrf1 α in protecting cells against cellular senescence induced by oxidative stress should be endowed with its unique capacity to maintain robust cellular homeostasis. This is supported by further experimental evidence revealing that Nrf1α deficiency led directly to diminished expression of its targets STAG2 and SMC3, thereby compromising chromosome stability and perturbing cellular homeostasis in *Nrf1α*^−/−^ cells. On this base, the heightened dysfunction of cellular autophagy caused by the loss of <u>Nrf1</u> is much likely to further exacerbate the cellular dyshomeostatic disturbance in *Nrf1α*^−/−^ cells. However, hyperactive Nrf2 accumulated in *Nrf1α*^−/−^ cells does not effectively counteract escalated disruption of the cellular homeostasis, though it can also exerts a limited compensatory response for such deficiency of Nrf1 α. Overall, deletion of Nrf1 α leads to a fatal disruption of the cellular homeostasis insomuch as to exacerbate the senescence process induced by oxidative stress.

### 4.1 The cytoprotective role of Nrf1 in the anti-senescent response to oxidative stress

Two distinct models of oxidative stress-induced cellular senescence cell were herein established, so as to give a better understanding of Nrf1’s role in anti-senescent protective response to oxidative stress. Of them, a simplified senescence model was established by allowing for growth of cancer cells (i.e., HepG2 and HeLa) to achieve a high level of confluence and undergo extended periods of incubation, so that they can lead to the exhaustion of essential nutrients, e.g., glucose, serum, and glutamine from the surrounding medium, but also concurrently to give rise to elevated concentrations of metabolic byproducts, such as lactate, ammonium ions, and pro-inflammatory cytokines, which are thought to be toxic stress or even induce oxidative stress. Such scenario has the capacity to significantly affect cellular processes, potentially leading to cellular senescence [47–51]. The simply resulting senescence model is considered as a type of oxidative stress-induced cellular senescence model, which offers an advantage of being easily constructed. Besides, an additional D-gal-induced senescence model was also established insofar as to gain insights into pathological manifestations of oxidative stress-induced senescence. This is just attributable to the formation of advanced glycation end-products (AGEs) and also subsequent induction of oxidative stress by interaction of D-gal (as a naturally-occurring reducing sugar found within the human body) with free amines of peptide amino acids. The oxidative stress is induced by elevated concentrations of D-gal through an oxidative reaction to yield ROS, in which galactose is converted into aldose and hydroperoxide with the aid of galactose oxidase catalysis. Thereby, the D-gal-induced cell senescence model serves as a widely-used framework for studying senescence induced by oxidative stress, and is also frequently applied to explore preventive or therapeutic interventions for ageing [52, 53].

Oxidative stress-induced cellular senescence is currently recognized as a dynamic process [54], along with a series of interconnected steps with contextual logical connections, and is also accompanied by the evolution and diversification of the characteristics of senescent cells. Upon accumulation of intracellular ROS and free radicals, oxidative stress is initiated, leading to oxidative modifications and/or even damages to biological macromolecules, which, in turn, trigger another series of stress-defensing responses insomuch as to alter the cellular state. Beyond antioxidant and detoxifying responses driven by Nrf1/Nrf2-target genes, the activation of the DNA damage response (DDR) is also induced to result in the eventual arrest of the cell cycle. Thus, it is, in the initiation of cellular senescence, found that the cells are propelled into the early stage known as ‘primary senescence’. During this phase, the stressed cells are like to have the ability to avoid cell cycle cessation and also potentially reverse senescence by promptly addressing the cellular damage caused by free radicals and/or other reactive species (including ROS). However, elevated levels of oxidative stress and its prolonged exposure result in heightened and accumulated cellular damages, along with impaired repair mechanisms and worsened metabolic imbalances. Consequently, this intensifies cell cycle arrest, ultimately propelling cells towards the subsequent senescence phase referred to as ‘developing senescence’. If such conditions continue to trigger cellular senescence, the cells may advance to a later stage known as ‘late senescence’. During this phase, the cells display complete senescence characteristics, which are placed at an irreversible state of the cell-cycle arrest. The primary definitive feature of cellular senescence induced by oxidative stress is manifested by transformation of cell cycle arrest from a temporary transitional phase to another persistent stable condition, and also accompanied by the stress-induced dysregulation of various cell cycle regulatory molecules [55, 56]. Consistently, the prominent feature of loss of Nrf1’s function is evinced by such an enhanced cell cycle arrest of *Nrf1α*^−/−^ in two distinct cellular senescence models induced by oxidative stress. This mark indicates an important cytoprotective role of Nrf1 exerted in the anti-senescent response to oxidative stress.

### 4.2 The loss of Nrf1 exacerbates development of SASP by reprogramming metabolic signals

An increasing number of studies unravel that the progression of cellular senescence induced by oxidative stress is accompanied by a significant alteration in the evolution of SASP profiles based on intracellular metabolic reprogramming [57]. The stress-induced senescent cells have undergone alterations in the metabolic processes, including reduced ratio of NAD^+^/NADH, heightened glycolysis and lactic acid generation, augmented uptake of lipids, enhanced synthesis of fatty acids and cholesterol, elevated production of oxylipins, and accumulation of lipid droplets (which are essential for certain aspects of the SASP). Such reorganized metabolism (re)programming entails alterations in relevant metabolic signaling pathways within senescent cells, as characterized by perturbation of critical regulators, e.g., deactivation of p53 and PTEN (in this study and [30]), which initiates the response of downstream signaling axis, leading to induction of the PI3K-AKT and NF-kB signaling cascades [58, 59]. This distinctive reprogramming of cellular senescence metabolism to signaling responses offers the provision of energy and/or building blocks required for the synthesis and release of SASP factors. This is supported by further evidence showing that pyruvate dehydrogenase kinase 4 (PDK4) is involved in enhancing aerobic glycolysis of senescent cells and aiding in lactate production, and thereby, in turn, contributes to the generation of SASP through ROS-mediated mechanisms [60]. Furthermore, senescent cells exhibit increased levels of both lipid β-oxidation and fatty acid synthesis (e.g., by activation of the *FASN* gene) [61, 62], supporting the need for macromolecular building blocks and additional energy required for the SASP production. In senescent cells NF-kB and mTOR remain active to contribute to the maintenance of SASP through the enhancement of transcript stability and facilitation of translation processes [63–65]. Collectively, these demonstrate the metabolic signal reprogramming of senescent cells had laid a vital cornerstone for development of SASP [66].

A growing body of evidence reveals that Nrf1 plays a crucial role in regulating intracellular metabolism [29, 67, 68], because the loss of its function leads to marked metabolic signal reprogramming [67, 69–71]. In this process, Nrf1 exerts its unique indispensable functions for governing the intracellular redox and energy (including lipid, cholesterol, glucose) metabolic homeostasis and also maintaining protein homeostasis and even chromosomal stability [20, 22, 31]. A notable increase in glucose uptake in pancreatic islet β-cells results from decreased expression of Nrf1 [72]. This is also supported by our previous experimental findings [69, 71], revealing that the loss of Nrf1 led to heightened glucose absorption and glycolysis, elevated lactate levels, enhanced fatty acid production and lipid deposition, as well as increased synthesis and transportation of amino acids, particularly serine synthesis. Such metabolic reprogramming resulting from the loss of Nrf1 is accompanied by rearrangement of metabolism-relevant signaling pathways, such as sustained activation of both NF-kB and PI3K-Akt-mTOR signaling axes [69, 70], with suppressed expression of master regulators p53 and PTEN.

In this study on cellular senescence, we also observed that the loss of Nrf1α resulted in a notable decrease in the expression of p53 as a key regulator of model cell senescence induced by oxidative stress. Such loss of Nrf1 α also resulted, as expected, in an enhancement of the SASP development and the progression in the context of oxidative stress-induced cellular senescence. This proposition is evidenced by experiments unraveling that Nrf1 α deficiency leads to enhanced development of SASP in *Nrf1α*^−/−^ cells under oxidative stress-induced senescent conditions. This effect is manifested by examined indicators of the early SASP elements (e.g., IL-1α and IL-1β) and the late SASP components (e.g., IL-6, CXCL-3 and MMP1). The amplification of SASP by *Nrf1α*^−/−^ is not only evinced by a characteristic feature of heightened cellular senescence induced by oxidative stress caused by loss of Nrf1α, but also serves as a notable contributing factor to further deteriorate cellular senescence (and/or other processes). This is because various elements of the SASP possess the ability to uphold and strengthen the senescent state by further perpetuating and amplifying SASP signaling cascades through relevant autocrine signaling pathways [66, 73]. For example, IL-1α, as an early SASP factor, can sustain and also enhance the SASP signaling by positively modulating NF-kB activity through an autocrine feedforward mechanism. Consequently, this also establishes another favorable feedback loop with senescent triggers like ROS, thereby preserving and enhancing the senescent cell cycle arrest. Altogether, these demonstrate that deletion of Nrf1 substantially enhances the oxidative stress-induced cellular senescence, as evidenced by *Nrf1α*^−/−^ -leading SASP, that is a well-recognized marker of cellular senescence and also a crucial contributing element to the advancement of aging and age-related ailments.

### 4.3 Nrf1 acts as a direct regulator of STAG2 and SMC3 in the cohesin required for chromosomal homeostasis

Since cell homeostasis is a critical hallmark influencing the healthy longevity of cells, senescent cells resulting from oxidative stress are conversely characterized by disturbance of the cellular homeostasis due to progressive accumulation of oxidative damaged and/or impaired biomolecules [74]. Thus, sustenance of cellular homeostasis impacts the pace of oxidative stress-induced cellular senescence, and its effectiveness also depends predominantly on the mechanisms of chromosomal homeostasis maintenance, which entail the collaboration between chromosomal stability and resistance to oxidative stress, together with chromosomal repair, elimination of impaired chromosomal material, and chromosomal surveillance [7, 75]. Heretofore, the accumulating evidence revealed that Nrf1 is essentially required for maintaining the robust cellular homeostasis in various aspects, including metabolic regulation, proteasomal activity, redox balance, and genetic stability [22, 31, 38, 68, 69].

In this study, we discovered that Nrf1 is a direct regulator of STAG2 and SMC3, as two critical elements required for maintaining the chromosomal homeostasis and genetic stability. The essential core protein STAG2 and the ATPase subunit SMC3 (and SMC1, together with RAD21) constitute the universally expressed and evolutionary conserved multisubunit protein complex, called cohesin, which encircles chromatin [43]. Cohesin is known to have certain pivotal effects in preserving the structural and functional integrity of genetic materials, primarily ensuring sister chromatid cohesion and high-fidelity chromosome separation so as to facilitate proper chromosome aggregation, maintaining a proper concentration to facilitate the error-free DNA damage repair, and organizing the genome into large loops, such as topologically associated domains (TADs), in order to generate functional domains and thus regulate gene expression by integrating transcriptional factors, target gene promoters and enhancers, with RNA polymerase II [76, 77]. Besides, the effects of cohesin can also be exerted in facilitating specific patterns of alternative splicing of the indicated transcripts to modulate diversity of those gene expression profiles [78]. Upon mutually exclusive mutations of STAG2 and/or SMC3 in the complex, this leads inevitably to the cohesin dysfunction and ensuing chromosomal dyshomeostatic imbalances, including aneuploidy, DNA damages, gene deregulation and other abnormalities, ultimately resulting in the disruption of cellular homeostasis and hence triggering cell senescence or even death [79, 80].

The evidence has been provided in our present study, unraveling that the transcriptional expression of STAG2 and SMC3 is specifically regulated by Nrf1, rather than Nrf2, directly through the consensus *AREs* existing in their gene promoter regions. Further experimental evidence demonstrates that the loss of such Nrf1’s function indeed exacerbated the cellular senescence by *Nrf1α*^−/−^-disrupting the basal expression of STAG2 and SMC3, particularly in oxidative stress-induced senescence conditions. Taken together, it is inferable that deletion of Nrf1α is much likely to result in a dramatic disturbance in the chromosomal stability through substantial down-regulation of STAG2 and SMC3, consequently leading to cell dyshomeostasis, all of which deteriorate oxidative stress-induced cellular senescence.

### 4.4 The cellular senescence exacerbated by *Nrf1α*^−/−^-leading dysfunction of cell autophagy

Since as an internal catabolic process widely preserved across different organisms, cell autophagy functions as a mechanism for the proper quality control by promptly eliminating damaged, harmful, and unnecessary materials, and concurrently serves as a self-preservation strategy in response to detrimental stimuli (e.g., oxidative stress), such that it is intricately linked with cellular senescence [45, 81]. Herein, we found that loss of Nrf1 α leads to worsened dysfunction of cell autophagy caused by *Nrf1α*^−/−^-leading oxidative stress in two distinct cellular senescence models. The underlying molecular mechanisms are attributable to disrupting the core ATG complexes in *Nrf1α*^−/−^ cells, as evidenced by this study showing that the lack of Nrf1 α resulted in substantially down-regulated expression of all examined autophagy-related factors (ATG4A, ATG5, ATG7, ATG9A, ATG10, ATG12. ATG13, ULK1 and Beclin1). These data demonstrate that Nrf1 plays a key regulatory role in the expression of autophagy factors. This is also supported by another report that *ATG2A*, *ATG4D* and *ATG13* were upregulated by Nrf1 overexpression and hence exerted a vital cytoprotective effect [28]. Of note, the autophagy-associated receptors p62 and GABARAPL1 are identified as two direct targets of Nrf1, so as to regulate their transcriptional expression [82–84]. However, our evidence has been presented showing that abundances of LC3-II and p62 (at its protein, but not mRNA, levels) were markedly enhanced in *Nrf1α* ^−/−^ cells. Taken altogether, such accumulation of p62 and LC3-II is suggested to result from enhanced dysfunction of *Nrf1α*^−/−^-cell autophagy (to lysosomal degradation) in oxidative stress-induced senescence conditions.

Besides, the non-ATG proteins (including PTEN) are also effectively involved in managing dynamics of autophagy [85]. Such a meticulously controlled process thus depends on the activation of multiple signaling (e.g., PI3K-mTOR) networks. Our previous work demonstrated that the activation of the PI3K-AKT-MTOR pathway in *Nrf1α*^−/−^ cells [69] is attributable to inactivation of PTEN by the loss of this CNC-bZIP factor [30]. Hence, it is inferable that the loss of Nrf1α is much like to lead to the inhibition of autophagy by dysregulating multihierarchical autophagy signaling network, such as the PTEN-PI3K-AKT-mTOR signaling with other non-ATG proteins involved.

The antioxidant factor Nrf1 functions as an indispensable control of robust redox homeostasis so as to tightly govern intracellular ROS levels [38], in which exists an intricate relationship between ROS and autophagy regulated by Nrf1. Low amounts of ROS (to induce eustress) trigger autophagy to facilitate cellular repair and also uphold metabolic equilibrium, but elevated levels of ROS induce oxidative distress, which, in turn, initiates another distinctive autophagy accounting for a defensive mechanism against oxidative damage. The cell autophagy serves the dual purpose of eliminating oxidized macromolecules and controlling ROS production through the targeted removal of the primary ROS-generating organelles, mitochondria and peroxisomes, besides autophagy-lysosomal degradation pathway. In this process, it is required for the maintenance of the activity of those particular regulatory molecules (e.g., PTEN) to an active threshold. When levels of ROS accumulate to a high extent and exceed this critical threshold, the process of autophagy is inhibited. For instance, an overabundance of ROS results in the PTEN inactivation in *Nrf1α*^−/−^ cells, which leads to the PI3K-AKT-mTOR activation and ensuing suppression of cell autophagy. Such elevated ROS can also cause a reduction in cell autophagy by oxidizing those essential autophagy core proteins (e.g., ATG10) [86]. The disruption of autophagy can further initiate a detrimental feedforward loop, in which the failure to eliminate increased ROS does, conversely, results in an additional yield of ROS. This is supported by our prior work showing that the absence of Nrf1 leads to an overabundance of ROS within cells [38]. Here, we further reveals that intracellular ROS accumulation in *Nrf1α* ^−/−^ cells is exacerbated particularly during oxidative stress-induced cellular senescence. From such collective evidence, it is hypothesized that the loss of Nrf1α is much likely to exacerbate the detrimental feedback loop existing between elevated ROS levels and reduced autophagy.

In fact, cellular autophagy is also essential for maintaining robust cell homeostasis under normal and even stressful conditions, but its dysfunction may lead to certain pathological alterations, e.g., in the advancement of senescence [87]. This is substantiated by the hitherto accumulating evidence obtained from genetic and pharmacological experiments across various species, revealing that impaired autophagy contributes to the process of senescence. This is because the proper autophagy demonstrates an inverse relationship with age during healthy lifespan, but the impaired autophagy is contributable to accelerated senescence phenotype (in fibroblasts, muscle cells, and neurons). Such a compromised autophagy exacerbates the process of pathological aging and relevant diseases [88]; this is evidenced by experiments on long-lived mutant animals uncovering another positive correlation between enhanced autophagy and a deceleration in the aging process. Altogether, it is inferable that oxidative stress-induced cellular senescence is likely exacerbated by the loss of Nrf1 leading to dysfunction of autophagy through a plausible mechanism. Conversely, Nrf1 can employ a range of strategies to regulate the expression of core ATG proteins and their activity, and integrate multihierarchical signaling network with non-ATG proteins. These autophagy mechanisms are substantially impeded by *Nrf1α*^−/−^-enhanced oxidative stress, leading to disruption of cell homeostasis insomuch as to expedite the senescence progression. However, the intricate interplay between Nrf1 and autophagy in oxidative stress-induced senescence remains to be further studied.

### 4.5 Distinct contributions of Nrf1 and Nrf2 to mediating the anti-senescent response

Nrf1 and Nrf2 share homological structural characteristics, similar expression patterns, and identical DNA recognition sites and thus together participate in a myriad of cell processes [22]. However, their functions are not entirely analogous, but also distinctive or even opposite in some occasions, displaying intricate relationships [20]. Global knockout of Nrf1 in mice gives rise to embryonic lethality, primarily due to enhanced oxidative stress, inadequate erythropoiesis, and insufficient *globin* expression [89–91]. By contrast, knockout of Nrf2 does not result in developmental arrest in mice, but increases the susceptibility to chemical carcinogens and oxidative stress [92–94]. These underscore distinct bioactivity of between Nrf1 and Nrf2. When compared to the single knockout of Nrf1, double knockout of Nrf1 and Nrf2 leads to early embryonic lethality, with elevated ROS examined in mouse embryonic fibroblast cells [95, 96]. This suggests overlapping biological functions of Nrf1 and Nrf2 to coordinately mitigate both internal and external stressors, providing a robust multi-layered defensing mechanism against stress. This is supported by the evidence showing that forced high-expression of Nrf1 in neurons compensates for Nrf2 deficiency in maintaining cellular redox balance [97]. Moreover, overexpression of Nrf1 or Nrf2 can also effectively rescue erastin2-induced ferroptosis in either deficiency contexts [98], implying their mutual compensatory responses.

However, when compared to Nrf2, Nrf1 possesses distinct or unique physiological and biological functions in the cellular protection and stress resilience to greater extents, because its loss cannot be fully compensated by accumulated Nrf2 [22]. This is further validated by a recent report that Nrf1, rather than Nrf2, provides substantial cardioprotection against ischemic/reperfusion injury in adult mice through a unique dual mechanism involving proteasome activation and redox homeostatic balance [28]. With regard to equivalent pathways or identical targets, Nrf1 also exhibits more robust regulatory effects than Nrf2, albeit both factors positively regulates PGC1α so as to maintain mitochondrial integrity with normal functional homeostasis [38]. This is because marked down-regulation of PGC1α by *Nrf1α*^−/−^ cannot be rescued by overexpressed Nrf2, thereby leading to the decline of mitochondrial function. Similar results were obtained from this study revealing that hyperactive Nrf2 accumulated cannot also reverse *Nrf1α* ^−/−^-leading inhibition of RB phosphorylation, even though Nrf1 and Nrf2 are involved in protecting against oxidative stress-induced senescence by regulating their co-targets (e.g., RB). Of crucial importance, Nrf1, but not Nrf2, has its unique capacity to maintain chromosomal stability through transcriptional regulation of STAG2 and SMC3 within the cohesin. This accounts for a particular regulatory mechanism in maintaining chromosomal homeostasis and genome stability insomuch as to combat senescence induced by oxidative stress. Further experimental evidence has been provided in this study demonstrating that, although Nrf2 is required for anti-senescent response to oxidative stress, this effect is too limited to exert a full compensatory response for the loss of Nrf1. This is due to the fact that *Nrf1α*^−/−^-leading oxidative stress-induced senescence was not rescued by significantly accumulated Nrf2, but further execrated by silencing of *siNrf2* in the double deficient cells exhibiting typical features of senescence. From this, it is inferable that the abnormal buildup of Nrf2 in *Nrf1α*^−/−^ cells is still likely endowed to a certain extent of the cellular resistance to senescence.

Collectively, the essential role of Nrf1 in maintaining cellular homeostasis is central to resist against oxidative stress-induced senescence, whilst Nrf2 also concurrently operates, as the secondary defense mechanism, to protect cells in anti-senescence response to acute unforeseen oxidative stress. Upon exposure to oxidative stress, the loss of Nrf1’s function exacerbates the compromise of cellular antioxidant defenses, to give rise to heightened risks of cell senescence resulting from oxidative stress. Thereby, to achieve a ‘buying time’ for effective induction of Nrf1-mediated homeostatic response against senescence induced by oxidative stress, the cells should activate an acute compensatory mechanism involving the upregulation of Nrf2 in the emergence response. However, the extent of compensation for the initial anti-senescence response is rather constrained, because Nrf2 is incapable to completely replace Nrf1 in protecting against senescence induced by oxidative stress. This inadequacy results in a lasting impairment of antioxidant defenses due to the absence of Nrf1, leading to a persistent exacerbation of redox imbalance. The resulting disturbance in the cellular homeostasis is further deteriorated insomuch as to accelerate the senescence progression.

### 4.6 Concluding remarks

The hitherto accumulating evidence demonstrates that Nrf1 acts as a vital cornerstone of the cellular defense system in cytoprotecting against oxidative stress insofar as to maintain the redox homeostatic equilibrium. Further evidence has been provided in this study, supporting that Nrf1 exerts its essential function in protecting against cellular senescence induced by oxidative stress, hence highlighting this indispensable and irreplaceable role in maintaining cell homeostasis. Of importance, Nrf1 α is identified as a direct regulator of STAG2 and SMC3 in the cohesin required for chromosomal stability and homeostasis. Upon loss of Nrf1 α, oxidative stress-induced cellular senescence is exacerbated by *Nrf1α*^−/−^-disrupting cellular homeostasis, dysregulating cell autophagy, reprogramming metabolic signals towards its target genes, as evinced by typical SASP with relevant markers. Furthermore, we have also presented the evidence revealing distinct contributions of Nrf1 and Nrf2 to mediating anti-senescent responses to oxidative stress. Although Nrf2 is also required for defending against oxidative stress-induced cellular senescence, this effect is too limited to fully compensate for the loss of Nrf1’s function. As such being the case, *Nrf1α*^−/−^-leading oxidative stress-induced cellular senescence cannot be rescued by accumulated Nrf2, but is further deteriorated by silencing of Nrf2 in this double deficient cells. Thereby, it is inferable that such distinctive roles of Nrf1 and Nrf2 are much likely coordinately exerted in defending distinct stages of cellular senescence induced by different extents of oxidative stress. Overall, the living ‘fossil’-like Nrf1 should be similar to its ancient orthologues SKN-1 and CncC, which have the exclusive potential to function as a central regulator for anti-ageing process during their health lifespans. Therefore, this study also presents a great prospect for development of new strategies by precision targeting Nrf1 to promote healthy longevity but also delay ageing and related disorders.

## Supplementary Materials

The supporting information, including six supplemental figures and also one supplemental table.

## Data availability

All required data are presented in the paper and Supplementary Materials. The additional data supporting this study is available upon reasonable request from the corresponding author.

## Acknowledgements

We are grateful to all other colleagues in Zhang’s group for the supportive help and critical discussion for this study. This work was funded by the National Natural Science Foundation of China (NSFC with two projects 82073079 and 81872336) to Prof. Yiguo Zhang.

## Author contributions

D.L. designed and performed the experiments, collected all the relevant data, and wrote a draft of this manuscript with figures and supplemental information. M.W. did bioinformatics analysis and experimental technical assistance. L.Q. had done design and construction of relevant knockout cell lines. S.H. refined relevant experimental methods and technical support. Lastly, Y.Z. designed and supervised this study, discussed and analyzed all the data, helped to prepare all figures with cartoons, rewrote and revised the paper. All these co-authors have read and agreed to the published version of the manuscript.

## Supplemental information

**Fig. S1.**
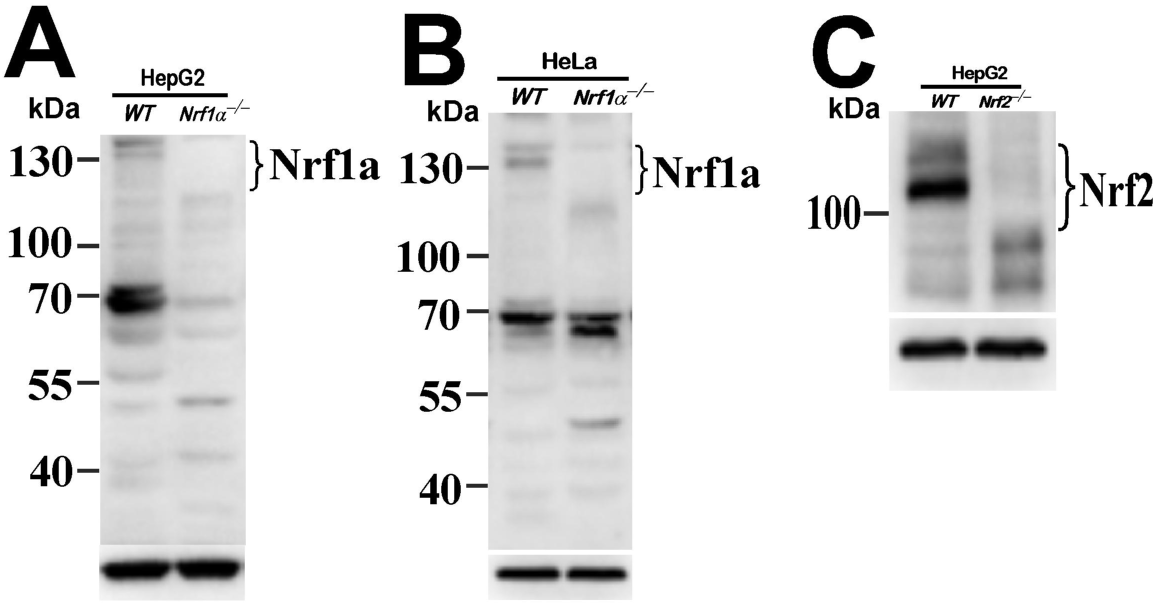
Identification of *Nrf1α*^−/−^ and *Nrf2*^−/−^ cell lines. (A) Distinction in the electrophoretic ability of Nrf1α between WT and Nrf1α^−/−^ derived from HepG2 cells as described previously. (B) The protein expression of Nrf1α in HeLa cells was completely abolished in Nrf1α^−/−^ cells. (C) Loss of Nrf2 was validated in Nrf2^−/−^ cells derived from HepG2 cells.

**Fig. S2.**
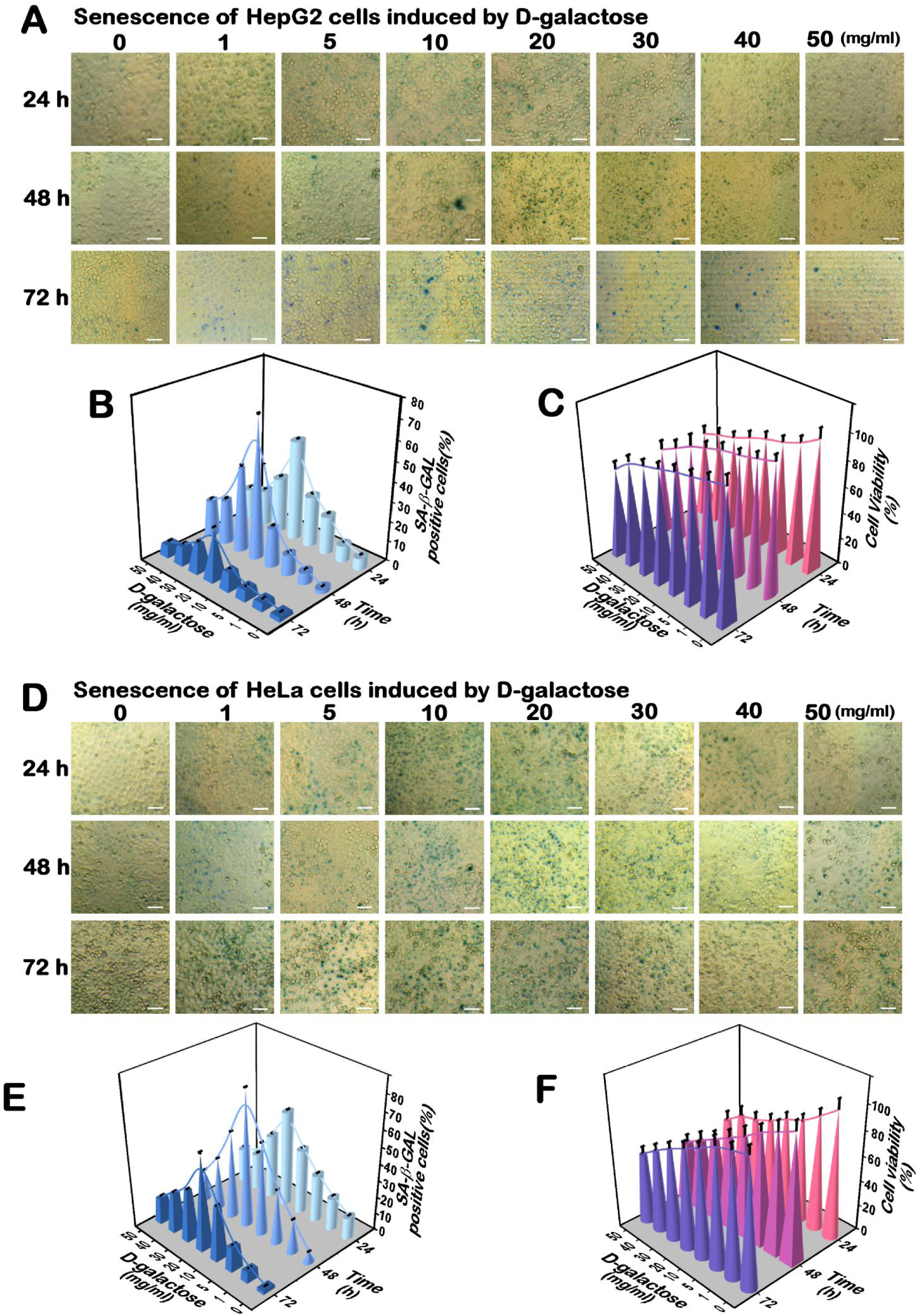
Two distinct cellular senescence models are established herein. (A-C) HepG2 cells were allowed for continuous culture for distinct periods of time (24, 48, or 72 h) and also subjected to treatment with distinct concentrates of D-gal (1, 5, 10, 20, 30, 40, and 50 mg/mL) for various durations (24, 48, or 72 h). The SA-β-gal positive staining cells are represented by relevant images (A, each bar = 50 μm), and the quantitative data are shown graphically as Mean values of at least three independent experiments performed in triplicates (B, n = 3 × 3). Subsequently, the above-described cell viability (C) was assayed by MTT. The data are shown as Mean ± SD, which were calculated from at least three independent experiments performed in triplicates (n = 3 × 3). (D-F) HeLa cells were allowed for continuous culture for distinct periods of time (24, 48, or 72 h) and also subjected to treatment with distinct concentrates of D-gal (1, 5, 10, 20, 30, 40, and 50 mg/mL) for various durations (24, 48, or 72 h). The SA-β-gal positive staining cells are represented by relevant images (D, each bar = 50 μm), and the quantitative data are shown graphically as Mean values of at least three independent experiments performed in triplicates (E, n = 3 × 3). Subsequently, the above-described cell viability (F) was assayed by MTT. The data are shown as Mean ± SD, which were calculated from at least three independent experiments performed in triplicates (n = 3 × 3).

**Fig. S3.**
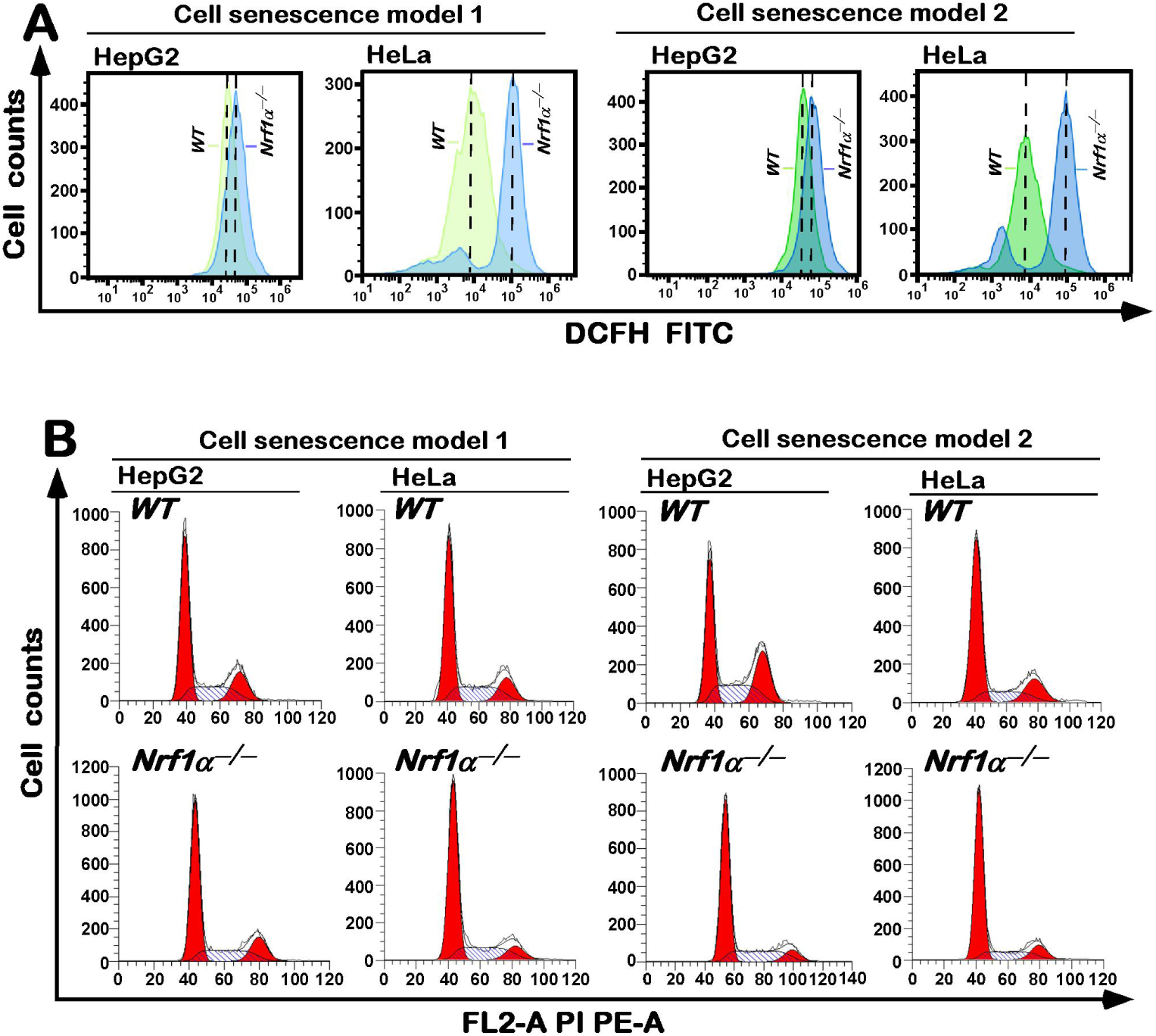
Changes in intracellular ROS levels and cell cycle of between *WT* and *Nrf1α*^−/−^ lines. (A) Flow cytometry analysis of intracellular ROS levels by using DCFH-DA fluorescent probes of *WT* and *Nrf1α*^−/−^ cell lines. (B) Distinct cell cycles between *WT* and *Nrf1α*^−/−^ lines were determined by flow cytometry and shown in original histograms.

**Fig. S4.**
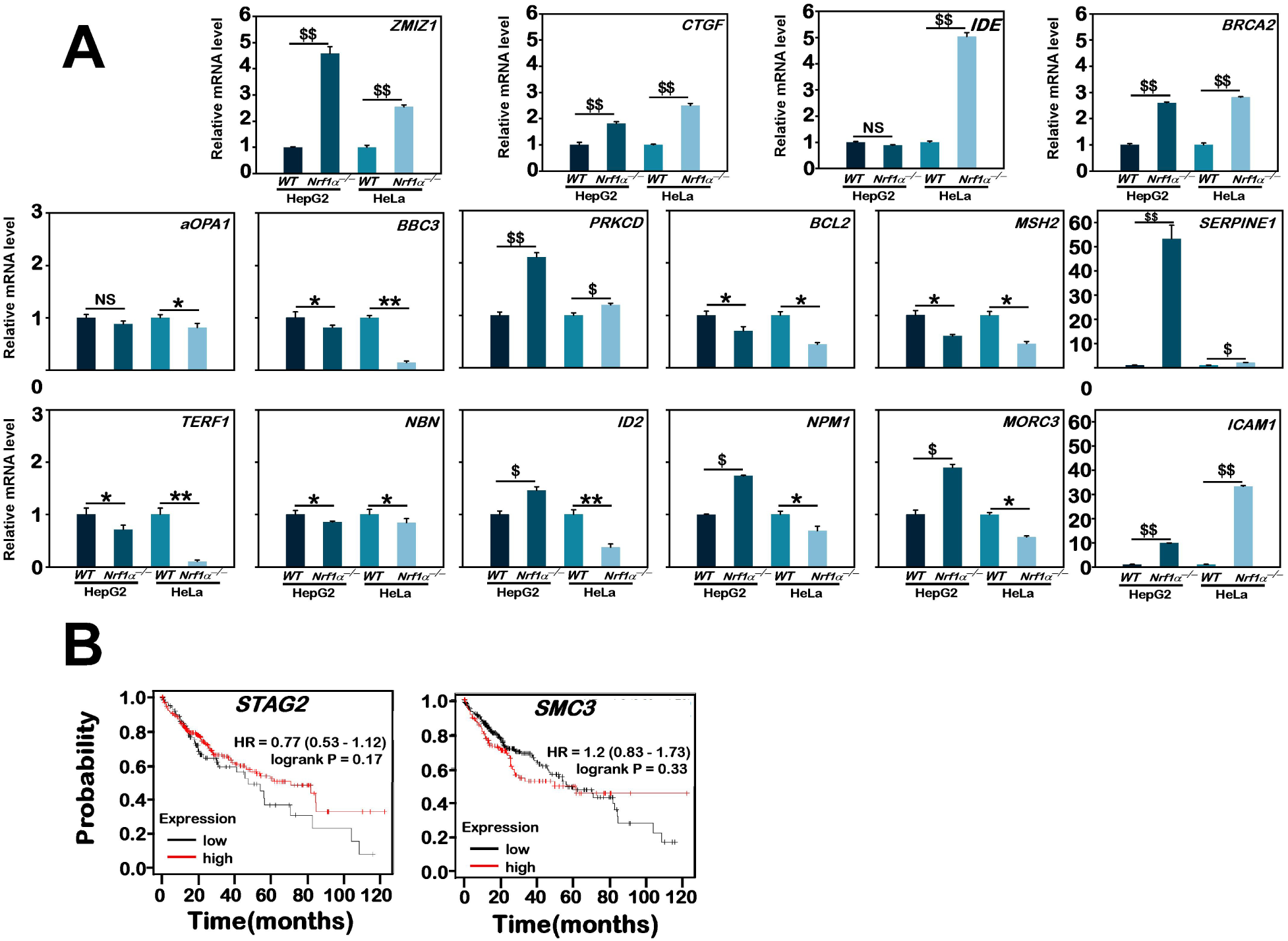
Those critical DEGs regulated by Nrf1 were further identified. (A) Distinct mRNA expression levels of selected key genes, *aOPA1, BBC3, PRKCD, BCL2, MSH2, TERF1, NBN, ID2, NPM1, MORC3, ZMIZ1, CTGF, IDE, BRCA2, ICAM1,* and *SERPINE1* in *WT* and *Nrf1α*^−/−^ cell lines were further validated by real-time qPCR. The resulting data are shown as Mean ± SD, with significant increases ($, p < 0.05; $$, p < 0.01), significant decreases (*, p < 0.05; **, p < 0.01;) or no statistical differences (NS), which were statistically determined from at least three independent experiments performed in triplicates (n = 3 × 3). (B) The correlation analysis of STAG2 and SMC3 with relevant OS rates of patients with HCC. The data were obtained from the online Kaplan-Meier plotter database.

**Fig. S5.**
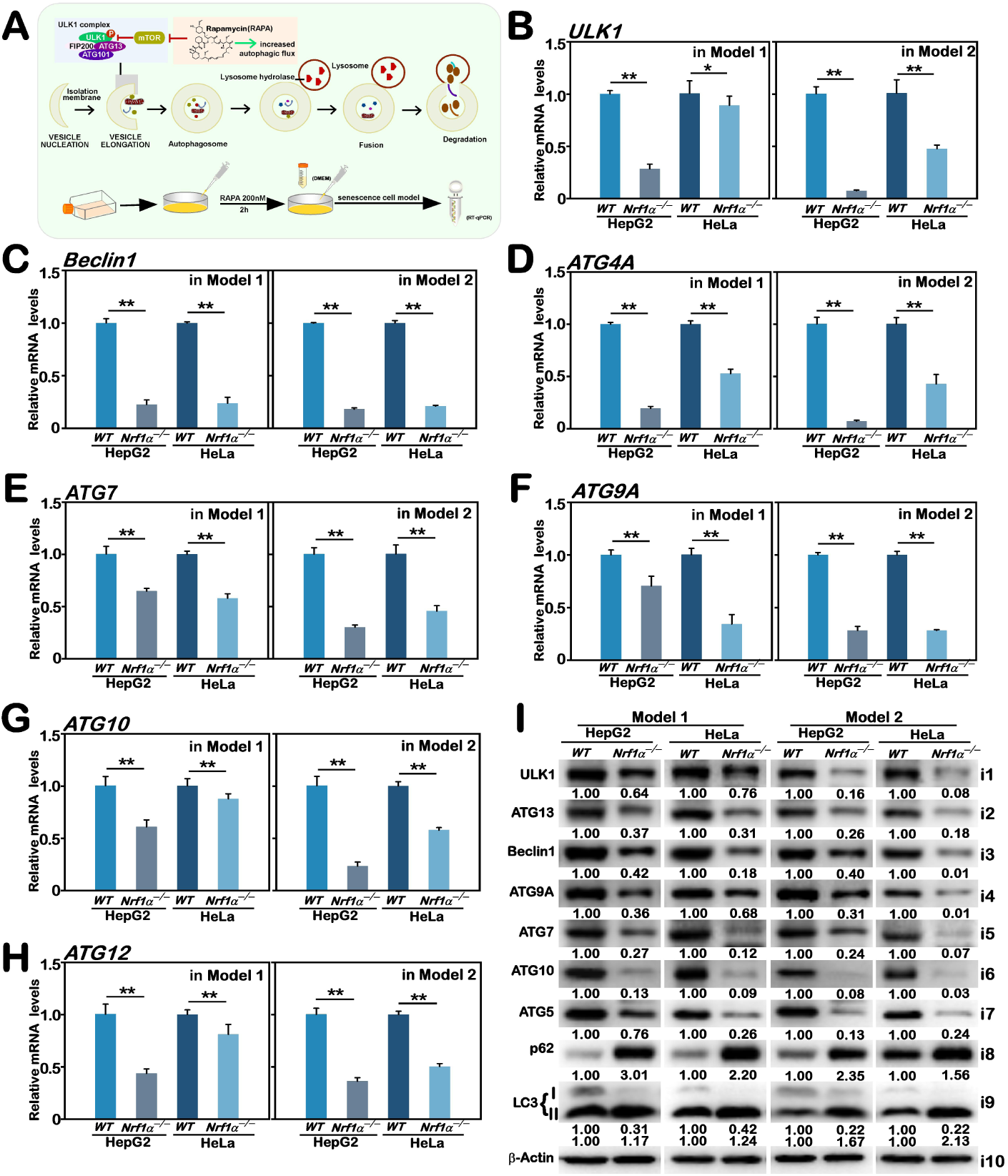
Potential effects of *Nrf1α*^−/−^-deficiency on the autophagy flux of cells treated with Rapamycin. (A) A schematic shows changes in the flux of cell autophagy stimulated by 200 nM of Rapamycin. (B–H) Real-time qPCR analysis of altered mRNA expression levels of autophagy-related markers, *ULK1* (B)*, Beclin1* (C)*, ATG4A*(D)*, ATG7* (E)*, ATG9A* (F)*, ATG10* (G), and *ATG12* (H) in *WT* and *Nrf1α*^−/−^ cell lines under distinct model senescence conditions. The resulting data are shown as Mean ± SD, with significant decreases (*, p < 0.05; **, p < 0.01), which were statistically determined from at least three independent experiments performed in triplicates (n = 3 × 3). (I) Western blotting analysis of altered protein expression levels of ULK1, Beclin1, ATG5, ATG7, ATG9A, ATG10, ATG13, p62, and LC3 in *WT* and *Nrf1α*^−/−^ cell lines under distinct model senescence conditions. The intensity of all immunoblots was quantified and then normalized by that of β-actin, before being shown as fold changes (*on the bottom* of indicated blots). The data presented herein were a representative of three independent experiments.

**Fig. S6.**
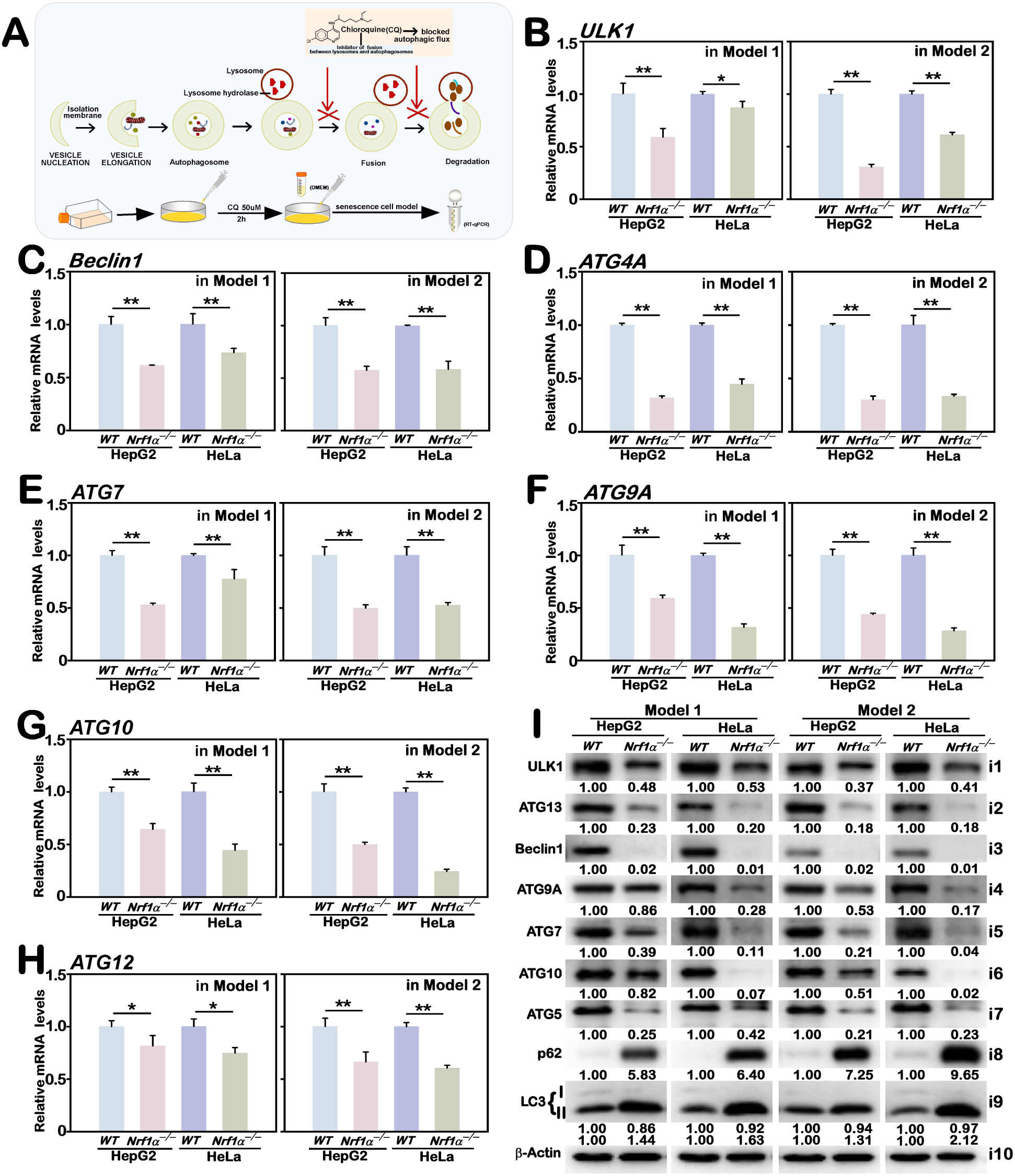
Potential effects of *Nrf1α*^−/−^-deficiency on the autophagy flux of cells treated with chloroquine. (A) A schematic shows changes in the flux of cell autophagy stimulated by 50 µM of chloroquine. (B–H) Real-time qPCR analysis of altered mRNA expression levels of autophagy-related markers, *ULK1* (B)*, Beclin1* (C)*, ATG4A*(D)*, ATG7* (E)*, ATG9A* (F)*, ATG10* (G), and *ATG12* (H) in *WT* and *Nrf1α*^−/−^ cell lines under distinct model senescence conditions. The resulting data are shown as Mean ± SD, with significant decreases (*, p < 0.05; **, p < 0.01), which were statistically determined from at least three independent experiments performed in triplicates (n = 3 × 3). (I) Western blotting analysis of altered protein expression levels of ULK1, Beclin1, ATG5, ATG7, ATG9A, ATG10, ATG13, p62, and LC3 in *WT* and *Nrf1α*^−/−^ cell lines under distinct model senescence conditions. The intensity of all immunoblots was quantified and then normalized by that of β-actin, before being shown as fold changes (*on the bottom* of indicated blots). The data presented herein were a representative of three independent experiments.

**Table S1.**
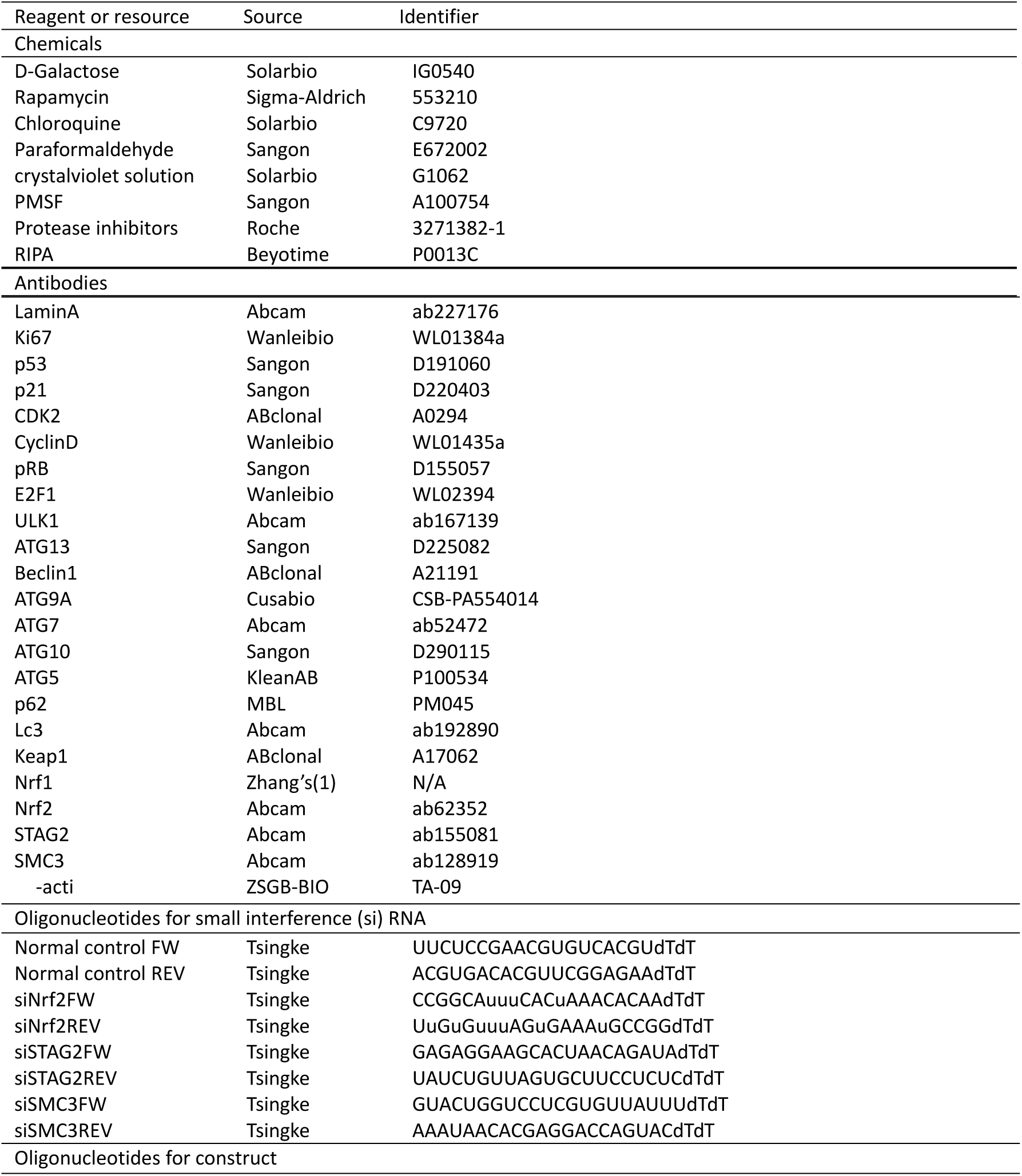
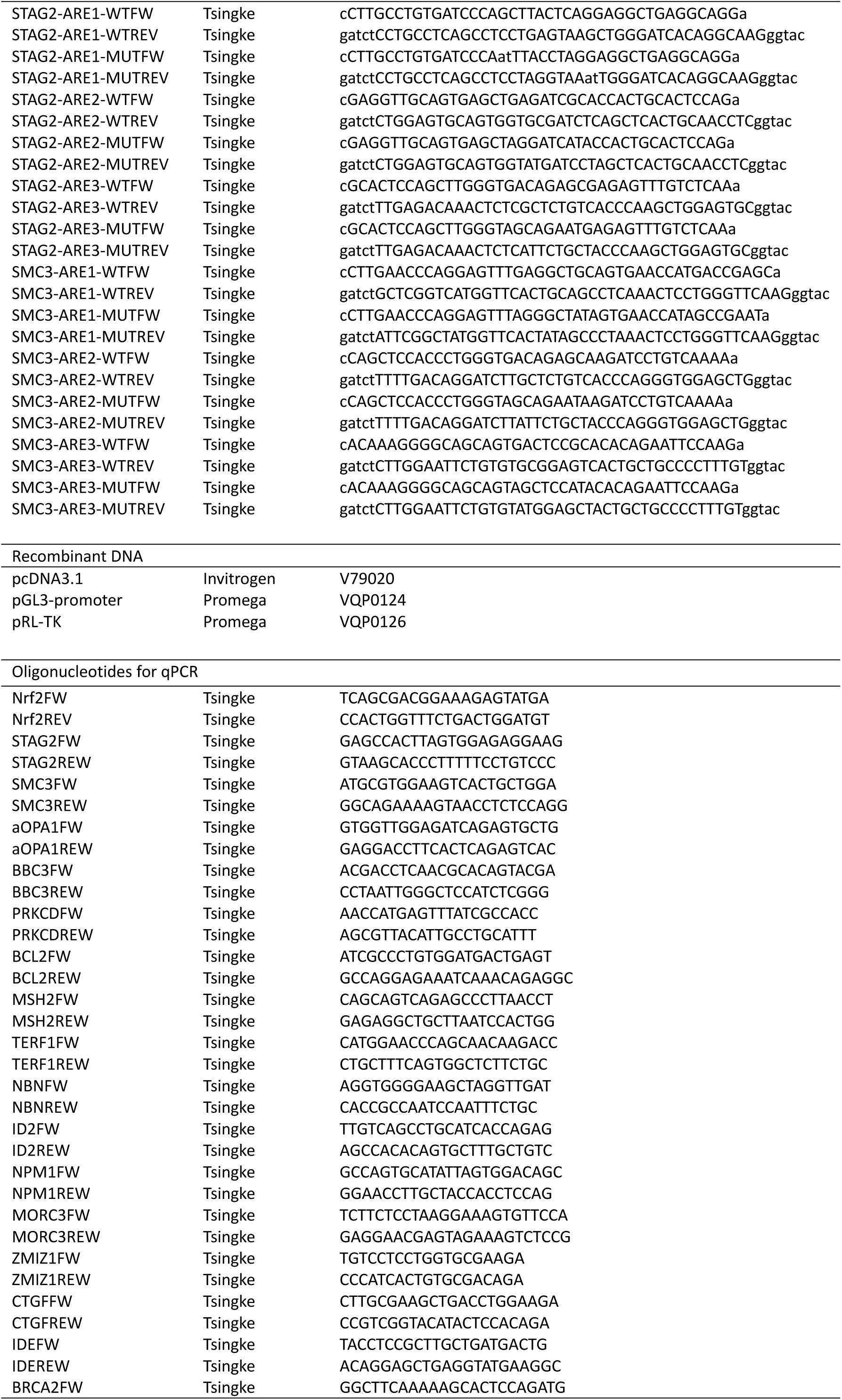
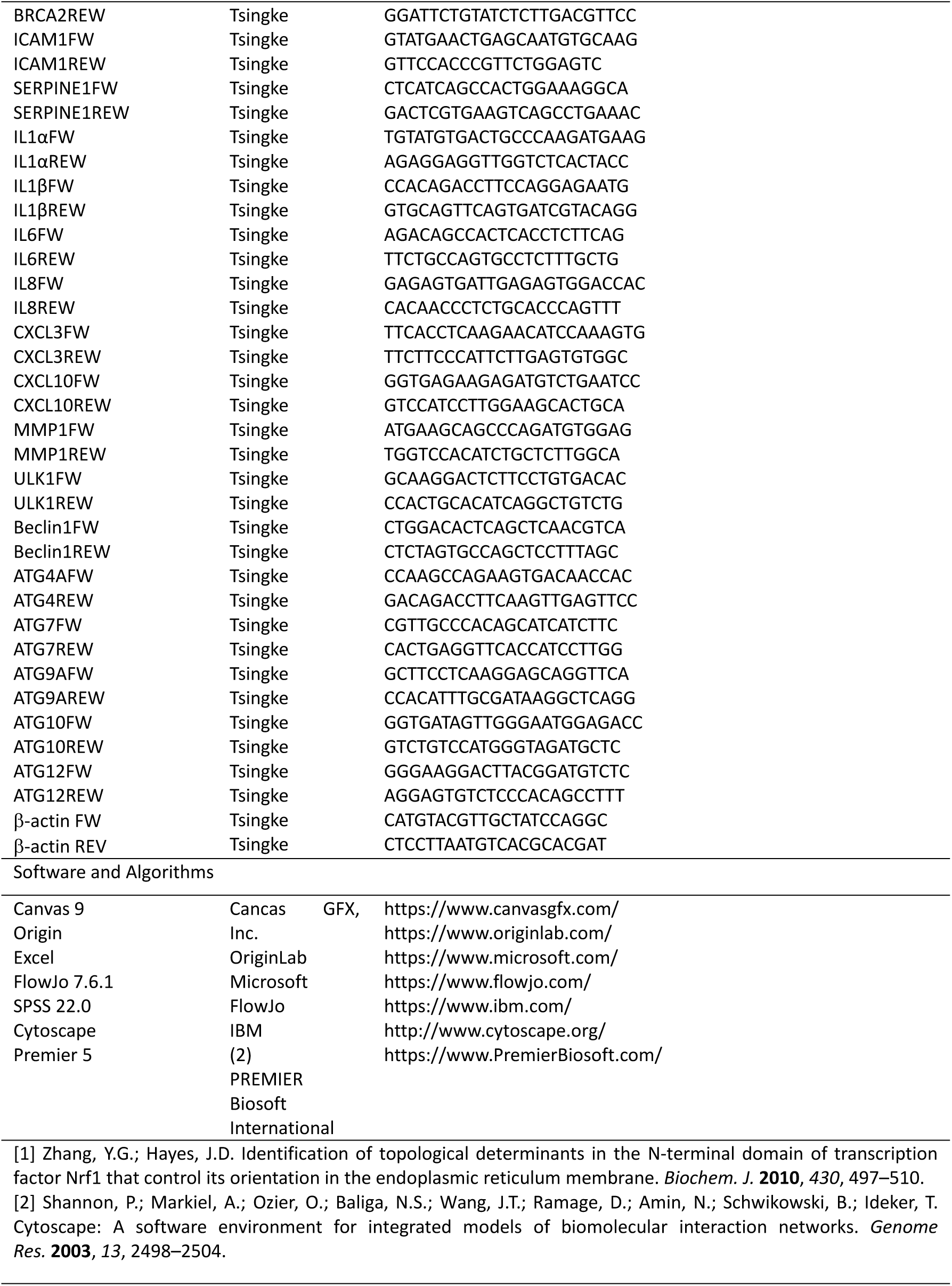
The key materials and relevant resources used in this work.

## References

1. Panyard, D. J., Yu, B. & Snyder, M. P. (2022) The metabolomics of human aging: Advances, challenges, and opportunities, Science Advances. 8, eadd6155.

2. Capitano, M. L., Mohamad, S. F., Cooper, S., Guo, B., Huang, X., Gunawan, A. M., Sampson, C., Ropa, J., Srour, E. F. & Orschell, C. M. (2021) Mitigating oxygen stress enhances aged mouse hematopoietic stem cell numbers and function, The Journal of clinical investigation. 131.

3. Schultz, M. B., Kane, A. E., Mitchell, S. J., MacArthur, M. R., Warner, E., Vogel, D. S., Mitchell, J. R., Howlett, S. E., Bonkowski, M. S. & Sinclair, D. A. (2020) Age and life expectancy clocks based on machine learning analysis of mouse frailty, Nature communications. 11, 4618.

4. Organization, W. H. (2022) Aging and health (2021) https://www.who.int/news-room/fact-sheets/detail/ageing-and-health, Accessed Jan. 29.

5. Lehallier, B., Gate, D., Schaum, N., Nanasi, T., Lee, S. E., Yousef, H., Moran Losada, P., Berdnik, D., Keller, A. & Verghese, J. (2019) Undulating changes in human plasma proteome profiles across the lifespan, Nature medicine. 25, 1843–1850.

6. Scott, A. J., Ellison, M. & Sinclair, D. A. (2021) The economic value of targeting aging, Nature Aging. 1, 616–623.

7. López-Otín, C., Blasco, M. A., Partridge, L., Serrano, M. & Kroemer, G. (2023) Hallmarks of aging: An expanding universe, Cell. 186, 243–278.

8. López-Otín, C., Pietrocola, F., Roiz-Valle, D., Galluzzi, L. & Kroemer, G. (2023) Meta-hallmarks of aging and cancer, Cell metabolism. 35, 12–35.

9. Guo, J., Huang, X., Dou, L., Yan, M., Shen, T., Tang, W. & Li, J. (2022) Aging and aging-related diseases: from molecular mechanisms to interventions and treatments, Signal Transduction and Targeted Therapy. 7, 391.

10. Avelar, R. A., Ortega, J. G., Tacutu, R., Tyler, E. J., Bennett, D., Binetti, P., Budovsky, A., Chatsirisupachai, K., Johnson, E. & Murray, A. (2020) A multidimensional systems biology analysis of cellular senescence in aging and disease, Genome biology. 21, 1–22.

11. Zhu, N., Liu, X., Xu, M. & Li, Y. (2021) Dietary nucleotides retard oxidative stress-induced senescence of human umbilical vein endothelial cells, Nutrients. 13, 3279.

12. Tchkonia, T., Palmer, A. K. & Kirkland, J. L. (2021) New horizons: novel approaches to enhance healthspan through targeting cellular senescence and related aging mechanisms, The Journal of Clinical Endocrinology & Metabolism. 106, e1481–e1487.

13. Wan, Y., Liu, J., Mai, Y., Hong, Y., Jia, Z., Tian, G., Liu, Y., Liang, H. & Liu, J. (2024) Current advances and future trends of hormesis in disease, npj Aging. 10, 26.

14. Shao, C.-S., Zhou, X.-H., Miao, Y.-H., Wang, P., Zhang, Q.-Q. & Huang, Q. (2021) In situ observation of mitochondrial biogenesis as the early event of apoptosis, Iscience. 24.

15. Jones, D. P. (2015) Redox theory of aging, Redox biology. 5, 71–79.

16. Ruvkun, G. & Lehrbach, N. (2023) Regulation and functions of the ER-associated Nrf1 transcription factor, Cold Spring Harbor Perspectives in Biology. 15, a041266.

17. Zhu, Y., Wang, M., Xiang, Y., Qiu, L., Hu, S., Zhang, Z., Mattjus, P. & Zhang, Y. (2018) Nach is a novel ancestral subfamily ofthe CNC-bZIP transcription factors selected during evolution from the marine bacteria to human, bioRxiv, 287755.

18. Zhang, H., Davies, K. J. & Forman, H. J. (2015) Oxidative stress response and Nrf2 signaling in aging, Free Radical Biology and Medicine. 88, 314–336.

19. Blackwell, T. K., Steinbaugh, M. J., Hourihan, J. M., Ewald, C. Y. & Isik, M. (2015) SKN-1/Nrf, stress responses, and aging in Caenorhabditis elegans, Free Radical Biology and Medicine. 88, 290–301.

20. Łuczyńska, K., Zhang, Z., Pietras, T., Zhang, Y. & Taniguchi, H. (2023) NFE2L1/Nrf1 serves as a potential therapeutical target for neurodegenerative diseases, Redox Biology, 103003.

21. Zhang, Y., Ren, Y., Li, S. & Hayes, J. D. (2014) Transcription factor Nrf1 is topologically repartitioned across membranes to enable target gene transactivation through its acidic glucose-responsive domains, PloS one. 9, e93458.

22. Zhang, Y. & Xiang, Y. (2016) Molecular and cellular basis for the unique functioning of Nrf1, an indispensable transcription factor for maintaining cell homoeostasis and organ integrity, Biochem J. 473, 961–1000.

23. Steffen, J., Seeger, M., Koch, A. & Krüger, E. (2010) Proteasomal degradation is transcriptionally controlled by TCF11 via an ERAD-dependent feedback loop, Molecular cell. 40, 147–158.

24. Xiang, Y., Wang, M., Hu, S., Qiu, L., Yang, F., Zhang, Z., Yu, S., Pi, J. & Zhang, Y. (2018) Mechanisms controlling the multistage post-translational processing of endogenous Nrf1α/TCF11 proteins to yield distinct isoforms within the coupled positive and negative feedback circuits, Toxicology and applied pharmacology. 360, 212–235.

25. Zhang, Y., Li, S., Xiang, Y., Qiu, L., Zhao, H. & Hayes, J. D. (2015) The selective post-translational processing of transcription factor Nrf1 yields distinct isoforms that dictate its ability to differentially regulate gene expression, Scientific reports. 5, 12983.

26. Kim, H. M., Han, J. W. & Chan, J. Y. (2016) Nuclear factor erythroid-2 like 1 (NFE2L1): structure, function and regulation, Gene. 584, 17–25.

27. Lehrbach, N. J. & Ruvkun, G. (2019) Endoplasmic reticulum-associated SKN-1A/Nrf1 mediates a cytoplasmic unfolded protein response and promotes longevity, Elife. 8, e44425.

28. Cui, M., Atmanli, A., Morales, M. G., Tan, W., Chen, K., Xiao, X., Xu, L., Liu, N., Bassel-Duby, R. & Olson, E. N. (2021) Nrf1 promotes heart regeneration and repair by regulating proteostasis and redox balance, Nature communications. 12, 5270.

29. Liu, X., Xu, C., Xiao, W. & Yan, N. (2023) Unravelling the role of NFE2L1 in stress responses and related diseases, Redox Biology, 102819.

30. Qiu, L., Wang, M., Hu, S., Ru, X., Ren, Y., Zhang, Z., Yu, S. & Zhang, Y. (2018) Oncogenic activation of Nrf2, though as a master antioxidant transcription factor, liberated by specific knockout of the full-length Nrf1α that acts as a dominant tumor repressor, Cancers. 10, 520.

31. Wufuer, R., Liu, K., Feng, J., Wang, M., Hu, S., Chen, F., Lin, S. & Zhang, Y. (2024) Distinct mechanisms by which Nrf1 and Nrf2 as drug targets contribute to the anticancer efficacy of cisplatin on hepatoma cells, Free Radical Biology and Medicine. 213, 488–511.

32. Wang, M., Ren, Y., Hu, S., Liu, K., Qiu, L. & Zhang, Y. (2021) TCF11 Has a Potent Tumor-Repressing Effect Than Its Prototypic Nrf1α by definition of both similar yet different regulatory profiles, with a striking disparity from Nrf2, Frontiers in oncology. 11, 707032.

33. Langmead, B., Trapnell, C., Pop, M. & Salzberg, S. L. (2009) Ultrafast and memory-efficient alignment of short DNA sequences to the human genome, Genome biology. 10, 1–10.

34. Kim, D., Langmead, B. & Salzberg, S. L. (2015) HISAT: a fast spliced aligner with low memory requirements, Nature methods. 12, 357–360.

35. Li, B. & Dewey, C. N. (2011) RSEM: accurate transcript quantification from RNA-Seq data with or without a reference genome, BMC bioinformatics. 12, 1–16.

36. Bader, G. D. & Hogue, C. W. (2003) An automated method for finding molecular complexes in large protein interaction networks, BMC bioinformatics. 4, 1–27.

37. Chin, C.-H., Chen, S.-H., Wu, H.-H., Ho, C.-W., Ko, M.-T. & Lin, C.-Y. (2014) cytoHubba: identifying hub objects and sub-networks from complex interactome, BMC systems biology. 8, 1–7.

38. Hu, S., Feng, J., Wang, M., Wufuer, R., Liu, K., Zhang, Z. & Zhang, Y. (2022) Nrf1 is an indispensable redox-determining factor for mitochondrial homeostasis by integrating multi-hierarchical regulatory networks, Redox Biology. 57, 102470.

39. Hernandez-Segura, A., Nehme, J. & Demaria, M. (2018) Hallmarks of cellular senescence, Trends in cell biology. 28, 436–453.

40. Tchkonia, T. & Kirkland, J. L. (2018) Aging, cell senescence, and chronic disease: emerging therapeutic strategies, Jama. 320, 1319–1320.

41. Garrido, A. M., Kaistha, A., Uryga, A. K., Oc, S., Foote, K., Shah, A., Finigan, A., Figg, N., Dobnikar, L. & Jørgensen, H. (2022) Efficacy and limitations of senolysis in atherosclerosis, Cardiovascular Research. 118, 1713–1727.

42. Van Deursen, J. M. (2014) The role of senescent cells in ageing, Nature. 509, 439–446.

43. Morales, C. & Losada, A. (2018) Establishing and dissolving cohesion during the vertebrate cell cycle, Current opinion in cell biology. 52, 51–57.

44. Ornatowski, W., Lu, Q., Yegambaram, M., Garcia, A. E., Zemskov, E. A., Maltepe, E., Fineman, J. R., Wang, T. & Black, S. M. (2020) Complex interplay between autophagy and oxidative stress in the development of pulmonary disease, Redox biology. 36, 101679.

45. Rajendran, P., Alzahrani, A. M., Hanieh, H. N., Kumar, S. A., Ben Ammar, R., Rengarajan, T. & Alhoot, M. A. (2019) Autophagy and senescence: A new insight in selected human diseases, Journal of cellular physiology. 234, 21485–21492.

46. Tabibzadeh, S. (2023) Role of autophagy in aging: The good, the bad, and the ugly, Aging Cell. 22, e13753.

47. Nobile, M. S., Votta, G., Palorini, R., Spolaor, S., De Vitto, H., Cazzaniga, P., Ricciardiello, F., Mauri, G., Alberghina, L. & Chiaradonna, F. (2020) Fuzzy modeling and global optimization to predict novel therapeutic targets in cancer cells, Bioinformatics. 36, 2181–2188.

48. White, E. Z., Pennant, N. M., Carter, J. R., Hawsawi, O., Odero-Marah, V. & Hinton, C. V. (2020) Serum deprivation initiates adaptation and survival to oxidative stress in prostate cancer cells, Scientific reports. 10, 12505.

49. Zhang, Y., Liu, L., Qi, Y., Lou, J., Chen, Y., Liu, C., Li, H., Chang, X., Hu, Z. & Li, Y. (2024) Lactic acid promotes nucleus pulposus cell senescence and corresponding intervertebral disc degeneration via interacting with Akt, Cellular and Molecular Life Sciences. 81, 1–25.

50. Linville, R. M., DeStefano, J. G., Sklar, M. B., Chu, C., Walczak, P. & Searson, P. C. (2020) Modeling hyperosmotic blood–brain barrier opening within human tissue-engineered in vitro brain microvessels, Journal of Cerebral Blood Flow & Metabolism. 40, 1517–1532.

51. Yang, K., Lu, R., Mei, J., Cao, K., Zeng, T., Hua, Y., Huang, X., Li, W. & Yin, Y. (2024) The war between the immune system and the tumor-using immune biomarkers as tracers, Biomarker Research. 12, 51.

52. Azman, K. F. & Zakaria, R. (2019) D-Galactose-induced accelerated aging model: an overview, Biogerontology. 20, 763–782.

53. Li, X., Li, C., Zhang, W., Wang, Y., Qian, P. & Huang, H. (2023) Inflammation and aging: signaling pathways and intervention therapies, Signal Transduction and Targeted Therapy. 8, 239.

54. Sun, R., Feng, J. & Wang, J. (2024) Underlying mechanisms and treatment of cellular senescence-induced biological barrier interruption and related diseases, Aging and Disease. 15, 612.

55. Liao, Z., Yeo, H. L., Wong, S. W. & Zhao, Y. (2021) Cellular senescence: mechanisms and therapeutic potential, Biomedicines. 9, 1769.

56. Chini, C. C. S., Cordeiro, H. S., Tran, N. L. K. & Chini, E. N. (2024) NAD metabolism: Role in senescence regulation and aging, Aging cell. 23, e13920.

57. Prašnikar, E., Borišek, J. & Perdih, A. (2021) Senescent cells as promising targets to tackle age-related diseases, Ageing Research Reviews. 66, 101251.

58. Wiley, C. D. & Campisi, J. (2021) The metabolic roots of senescence: mechanisms and opportunities for intervention, Nature Metabolism. 3, 1290–1301.

59. Kim, Y., Jang, Y., Kim, M.-S. & Kang, C. (2024) Metabolic remodeling in cancer and senescence and its therapeutic implications, Trends in Endocrinology & Metabolism.

60. Ma, Z., Ding, Y., Ding, X., Mou, H., Mo, R. & Tan, Q. (2023) PDK4 rescues high-glucose-induced senescent fibroblasts and promotes diabetic wound healing through enhancing glycolysis and regulating YAP and JNK pathway, Cell Death Discovery. 9, 424.

61. Nehlin, J. O., Just, M., Rustan, A. C. & Gaster, M. (2011) Human myotubes from myoblast cultures undergoing senescence exhibit defects in glucose and lipid metabolism, Biogerontology. 12, 349–365.

62. Jung, Y. H., Chae, C. W., Chang, H. S., Choi, G. E., Lee, H. J. & Han, H. J. (2022) Silencing SIRT5 induces the senescence of UCB-MSCs exposed to TNF-α by reduction of fatty acid β-oxidation and anti-oxidation, Free Radical Biology and Medicine. 192, 1–12.

63. Kanigur Sultuybek, G., Soydas, T. & Yenmis, G. (2019) NF-κB as the mediator of metformin’s effect on ageing and ageing-related diseases, Clinical and Experimental Pharmacology and Physiology. 46, 413–422.

64. Saoudaoui, S., Bernard, M., Cardin, G. B., Malaquin, N., Christopoulos, A. & Rodier, F. (2021) mTOR as a senescence manipulation target: A forked road, Advances in cancer research. 150, 335–363.

65. Weichhart, T. (2018) mTOR as regulator of lifespan, aging, and cellular senescence: a mini-review, Gerontology. 64, 127–134.

66. Wang, B., Han, J., Elisseeff, J. H. & Demaria, M. (2024) The senescence-associated secretory phenotype and its physiological and pathological implications, Nat Rev Mol Cell Bio, 1–21.

67. Qiu, L., Yang, Q., Zhao, W., Xing, Y., Li, P., Zhou, X., Ning, H., Shi, R., Gou, S. & Chen, Y. (2022) Dysfunction of the energy sensor NFE2L1 triggers uncontrollable AMPK signaling and glucose metabolism reprogramming, Cell death & disease. 13, 501.

68. Ren, S., Bian, Y., Hou, Y., Wang, Z., Zuo, Z., Liu, Z., Teng, Y., Fu, J., Wang, H. & Xu, Y. (2021) The roles of NFE2L1 in adipocytes: Structural and mechanistic insight from cell and mouse models, Redox biology. 44, 102015.

69. Deng, R., Zhu, Y., Liu, K., Zhang, Q., Hu, S., Wang, M. & Zhang, Y. (2024) Genetic loss of Nrf1 and Nrf2 leads to distinct metabolism reprogramming of HepG2 cells by opposing regulation of the PI3K-AKT-mTOR signalling pathway, Bioorganic Chemistry, 107212.

70. Deng, R., Zheng, Z., Hu, S., Wang, M., Feng, J., Mattjus, P., Zhang, Z. & Zhang, Y. (2024) Loss of Nrf1 rather than Nrf2 leads to inflammatory accumulation of lipids and reactive oxygen species in human hepatoma cells, which is alleviated by 2-bromopalmitate, Biochimica et Biophysica Acta (BBA)-Molecular Cell Research. 1871, 119644.

71. Zhu, Y.-p., Zheng, Z., Xiang, Y. & Zhang, Y. (2020) Glucose Starvation-Induced Rapid Death of Nrf1α-Deficient, but Not Nrf2-Deficient, Hepatoma Cells Results from Its Fatal Defects in the Redox Metabolism Reprogramming, Oxidative medicine and cellular longevity. 2020, 4959821.

72. Zheng, H., Fu, J., Xue, P., Zhao, R., Dong, J., Liu, D., Yamamoto, M., Tong, Q., Teng, W. & Qu, W. (2015) CNC-bZIP protein Nrf1-dependent regulation of glucose-stimulated insulin secretion, Antioxidants & redox signaling. 22, 819–831.

73. Serrano, M. & Munoz-Espin, D. (2021) Cellular Senescence in Disease, Academic Press.

74. Yamagata, Y., Fukuyama, T., Onami, S. & Masuya, H. (2024) Prototyping an Ontological Framework for Cellular Senescence Mechanisms: A Homeostasis Imbalance Perspective, Scientific Data. 11, 485.

75. Suganuma, T. & Workman, J. L. (2023) Chromatin balances cell redox and energy homeostasis, Epigenetics & Chromatin. 16, 46.

76. Marko, J. F., De Los Rios, P., Barducci, A. & Gruber, S. (2019) DNA-segment-capture model for loop extrusion by structural maintenance of chromosome (SMC) protein complexes, Nucleic acids research. 47, 6956–6972.

77. Meisenberg, C., Pinder, S. I., Hopkins, S. R., Wooller, S. K., Benstead-Hume, G., Pearl, F. M., Jeggo, P. A. & Downs, J. A. (2019) Repression of transcription at DNA breaks requires cohesin throughout interphase and prevents genome instability, Molecular cell. 73, 212–223. e7.

78. Singh, A. K., Chen, Q., Nguyen, C., Meerzaman, D. & Singer, D. S. (2023) Cohesin regulates alternative splicing, Science Advances. 9, eade3876.

79. Cuartero, S., Weiss, F. D., Dharmalingam, G., Guo, Y., Ing-Simmons, E., Masella, S., Robles-Rebollo, I., Xiao, X., Wang, Y.-F. & Barozzi, I. (2018) Control of inducible gene expression links cohesin to hematopoietic progenitor self-renewal and differentiation, Nature immunology. 19, 932–941.

80. Pati, D. (2024) Role of chromosomal cohesion and separation in aneuploidy and tumorigenesis, Cellular and Molecular Life Sciences. 81, 100.

81. Revuelta, M. & Matheu, A. (2017) Autophagy in stem cell aging, Aging cell. 16, 912–915.

82. Hatanaka, A., Nakada, S., Matsumoto, G., Satoh, K., Aketa, I., Watanabe, A., Hirakawa, T., Tsujita, T., Waku, T. & Kobayashi, A. (2023) The transcription factor NRF1 (NFE2L1) activates aggrephagy by inducing p62 and GABARAPL1 after proteasome inhibition to maintain proteostasis, Scientific Reports. 13, 14405.

83. Sha, Z., Schnell, H. M., Ruoff, K. & Goldberg, A. (2018) Rapid induction of p62 and GABARAPL1 upon proteasome inhibition promotes survival before autophagy activation, Journal of Cell Biology. 217, 1757–1776.

84. Ward, M. A., Vangala, J. R., Kamber Kaya, H. E., Byers, H. A., Hosseini, N., Diaz, A., Cuervo, A. M., Kaushik, S. & Radhakrishnan, S. K. (2024) Transcription factor Nrf1 regulates proteotoxic stress-induced autophagy, Journal of Cell Biology. 223, e202306150.

85. Liang, S., Wu, Y.-S., Li, D.-Y., Tang, J.-X. & Liu, H.-F. (2022) Autophagy and renal fibrosis, Aging and disease. 13, 712.

86. Zhou, J., Li, X.-Y., Liu, Y.-J., Feng, J., Wu, Y., Shen, H.-M. & Lu, G.-D. (2022) Full-coverage regulations of autophagy by ROS: from induction to maturation, Autophagy. 18, 1240–1255.

87. Zhang, W., Nishimura, T., Gahlot, D., Saito, C., Davis, C., Jefferies, H. B., Schreiber, A., Thukral, L. & Tooze, S. A. (2023) Autophagosome membrane expansion is mediated by the N-terminus and cis-membrane association of human ATG8s, Elife. 12, e89185.

88. Aman, Y., Schmauck-Medina, T., Hansen, M., Morimoto, R. I., Simon, A. K., Bjedov, I., Palikaras, K., Simonsen, A., Johansen, T. & Tavernarakis, N. (2021) Autophagy in healthy aging and disease, Nature aging. 1, 634–650.

89. Farmer, S. C., Sun, C.-W., Winnier, G. E., Hogan, B. & Townes, T. M. (1997) The bZIP transcription factor LCR-F1 is essential for mesoderm formation in mouse development, Genes & development. 11, 786–798.

90. Chan, J. Y., Kwong, M., Lu, R., Chang, J., Wang, B., Yen, T. B. & Kan, Y. W. (1998) Targeted disruption of the ubiquitous CNC-bZIP transcription factor, Nrf-1, results in anemia and embryonic lethality in mice, The EMBO journal. 17, 1779–1787.

91. Chen, L., Kwong, M., Lu, R., Ginzinger, D., Lee, C., Leung, L. & Chan, J. Y. (2003) Nrf1 is critical for redox balance and survival of liver cells during development, Molecular and cellular biology. 23, 4673–4686.

92. Enomoto, A., Itoh, K., Nagayoshi, E., Haruta, J., Kimura, T., O’Connor, T., Harada, T. & Yamamoto, M. (2001) High sensitivity of Nrf2 knockout mice to acetaminophen hepatotoxicity associated with decreased expression of ARE-regulated drug metabolizing enzymes and antioxidant genes, Toxicol Sci. 59, 169–177.

93. Aoki, Y., Sato, H., Nishimura, N., Takahashi, S., Itoh, K. & Yamamoto, M. (2001) Accelerated DNA adduct formation in the lung of the Nrf2 knockout mouse exposed to diesel exhaust, Toxicology and applied pharmacology. 173, 154–160.

94. Xu, C., Huang, M.-T., Shen, G., Yuan, X., Lin, W., Khor, T. O., Conney, A. H. & Kong, A.-N. T. (2006) Inhibition of 7, 12-dimethylbenz (a) anthracene-induced skin tumorigenesis in C57BL/6 mice by sulforaphane is mediated by nuclear factor E2–related factor 2, Cancer research. 66, 8293–8296.

95. Leung, L., Kwong, M., Hou, S., Lee, C. & Chan, J. Y. (2003) Deficiency of the Nrf1 and Nrf2 transcription factors results in early embryonic lethality and severe oxidative stress, Journal of Biological Chemistry. 278, 48021–48029.

96. Sekine, H., Okazaki, K., Kato, K., Alam, M. M., Shima, H., Katsuoka, F., Tsujita, T., Suzuki, N., Kobayashi, A. & Igarashi, K. (2018) O-GlcNAcylation signal mediates proteasome inhibitor resistance in cancer cells by stabilizing NRF1, Molecular and Cellular Biology.

97. Lee, C. S., Lee, C., Hu, T., Nguyen, J. M., Zhang, J., Martin, M. V., Vawter, M. P., Huang, E. J. & Chan, J. Y. (2011) Loss of nuclear factor E2-related factor 1 in the brain leads to dysregulation of proteasome gene expression and neurodegeneration, Proceedings of the National Academy of Sciences. 108, 8408–8413.

98. Forcina, G. C., Pope, L., Murray, M., Dong, W., Abu-Remaileh, M., Bertozzi, C. R. & Dixon, S. J. (2022) Ferroptosis regulation by the NGLY1/NFE2L1 pathway, Proceedings of the National Academy of Sciences. 119, e2118646119.

