## Supplemental Figures and Table S1 for "Deletion of Nrf1 exacerbates oxidative stress-induced cellular senescence by disrupting the cell homeostasis"

Supplemental information

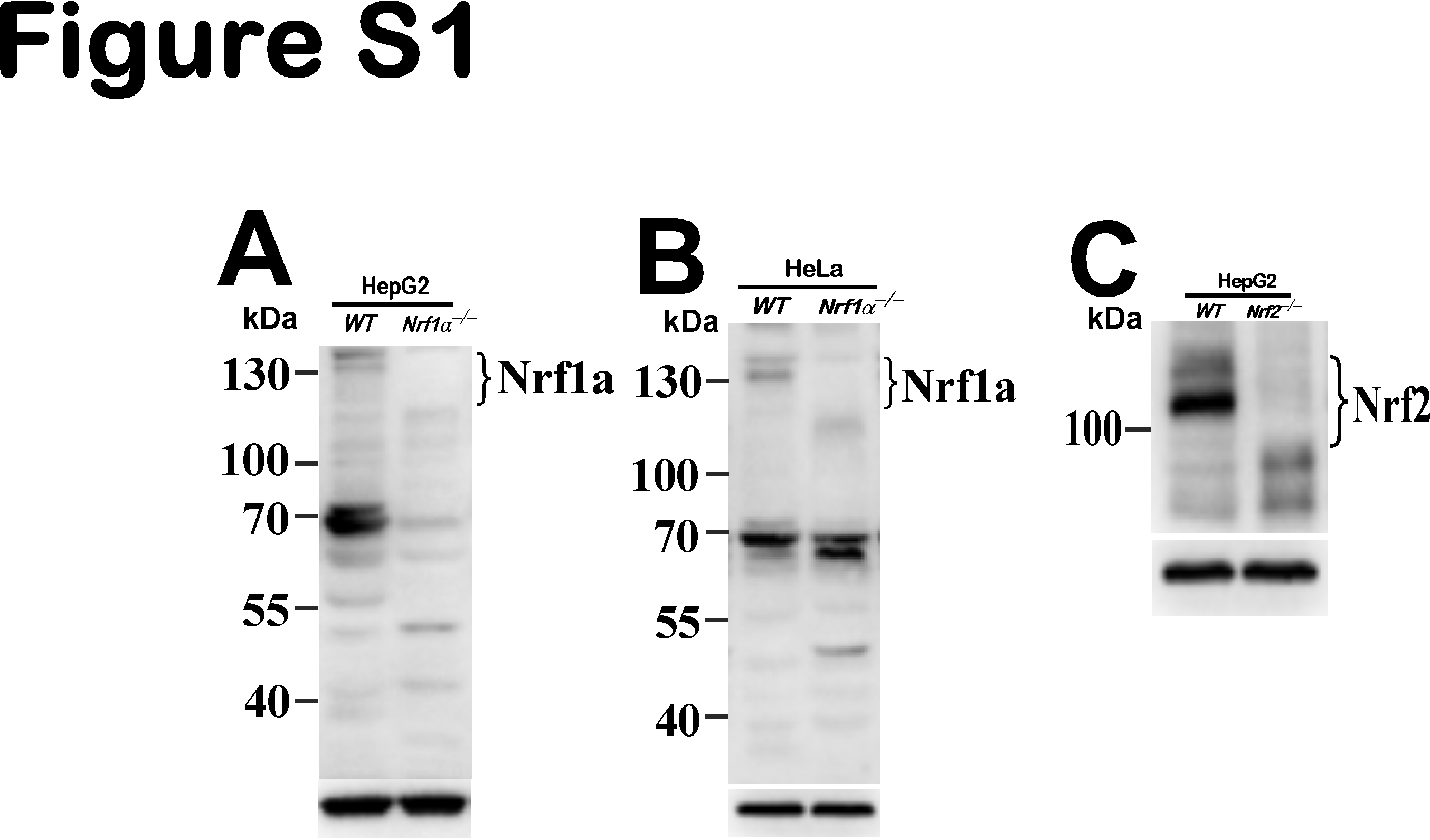

**Fig. S1. Identification of *Nrf1α^−/−^* and *Nrf2^−/−^* cell lines**

(A) Distinction in the electrophoretic ability of Nrf1α between WT and Nrf1α^−/−^ derived from HepG2 cells as described previously.

(B) The protein expression of Nrf1α in HeLa cells was completely abolished in Nrf1α^−/−^ cells.

(C) Loss of Nrf2 was validated in Nrf2^−/−^ cells derived from HepG2 cells.

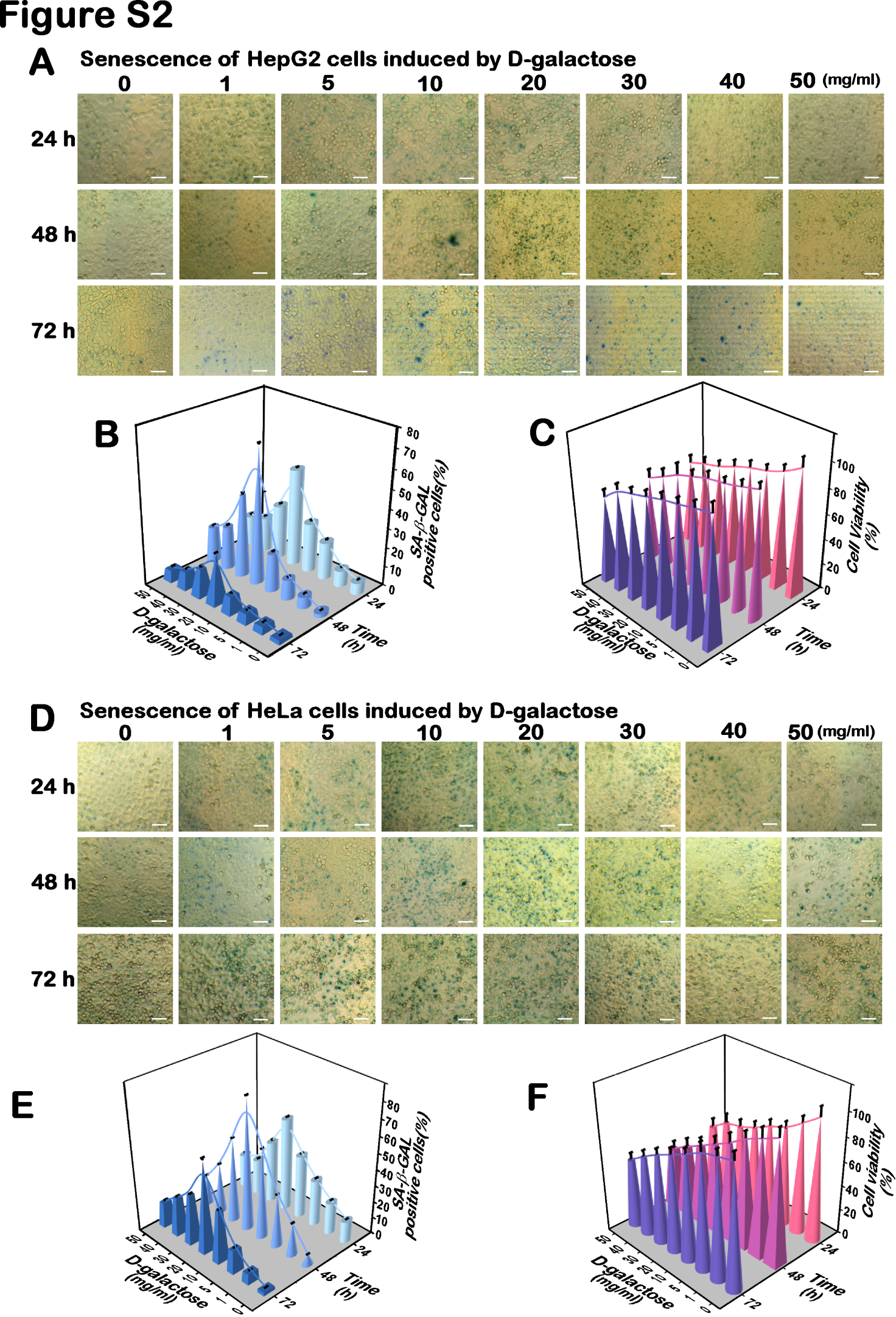

**Fig. S2. Two distinct cellular senescence models are** **established herein**

(A-C) HepG2 cells were allowed for continuous culture for distinct periods of time (24, 48, or 72 h) and also subjected to treatment with distinct concentrates of D-gal (1, 5, 10, 20, 30, 40, and 50 mg/mL) for various durations (24, 48, or 72 h). The SA-β-gal positive staining cells are represented by relevant images (A, each bar = 50 μm), and the quantitative data are shown graphically as Mean values of at least three independent experiments performed in triplicates (B, n = 3 × 3). Subsequently, the above-described cell viability (C) was assayed by MTT. The data are shown as Mean ± SD, which were calculated from at least three independent experiments performed in triplicates (n = 3 × 3).

(D-F) HeLa cells were allowed for continuous culture for distinct periods of time (24, 48, or 72 h) and also subjected to treatment with distinct concentrates of D-gal (1, 5, 10, 20, 30, 40, and 50 mg/mL) for various durations (24, 48, or 72 h). The SA-β-gal positive staining cells are represented by relevant images (D, each bar = 50 μm), and the quantitative data are shown graphically as Mean values of at least three independent experiments performed in triplicates (E, n = 3 × 3). Subsequently, the above-described cell viability (F) was assayed by MTT. The data are shown as Mean ± SD, which were calculated from at least three independent experiments performed in triplicates (n = 3 × 3).

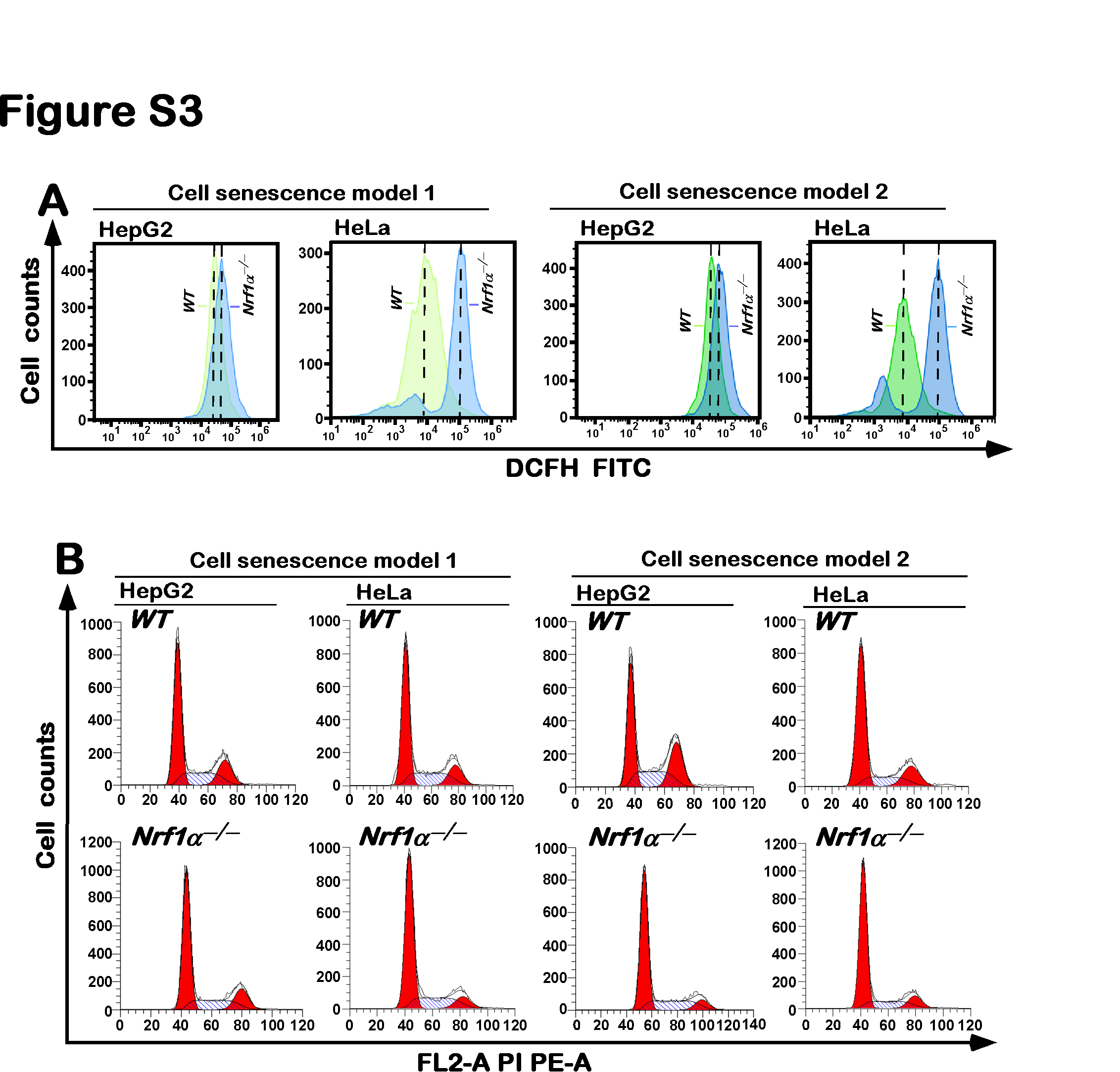

**Fig. S3. Changes in intracellular ROS levels and cell cycle of between *WT* and *Nrf1α^−/−^* lines**

(A) Flow cytometry analysis of intracellular ROS levels by using DCFH-DA fluorescent probes of *WT* and *Nrf1α^−/−^* cell lines.
(B) Distinct cell cycles between *WT* and *Nrf1α^−/−^* lines were determined by flow cytometry and shown in original histograms.

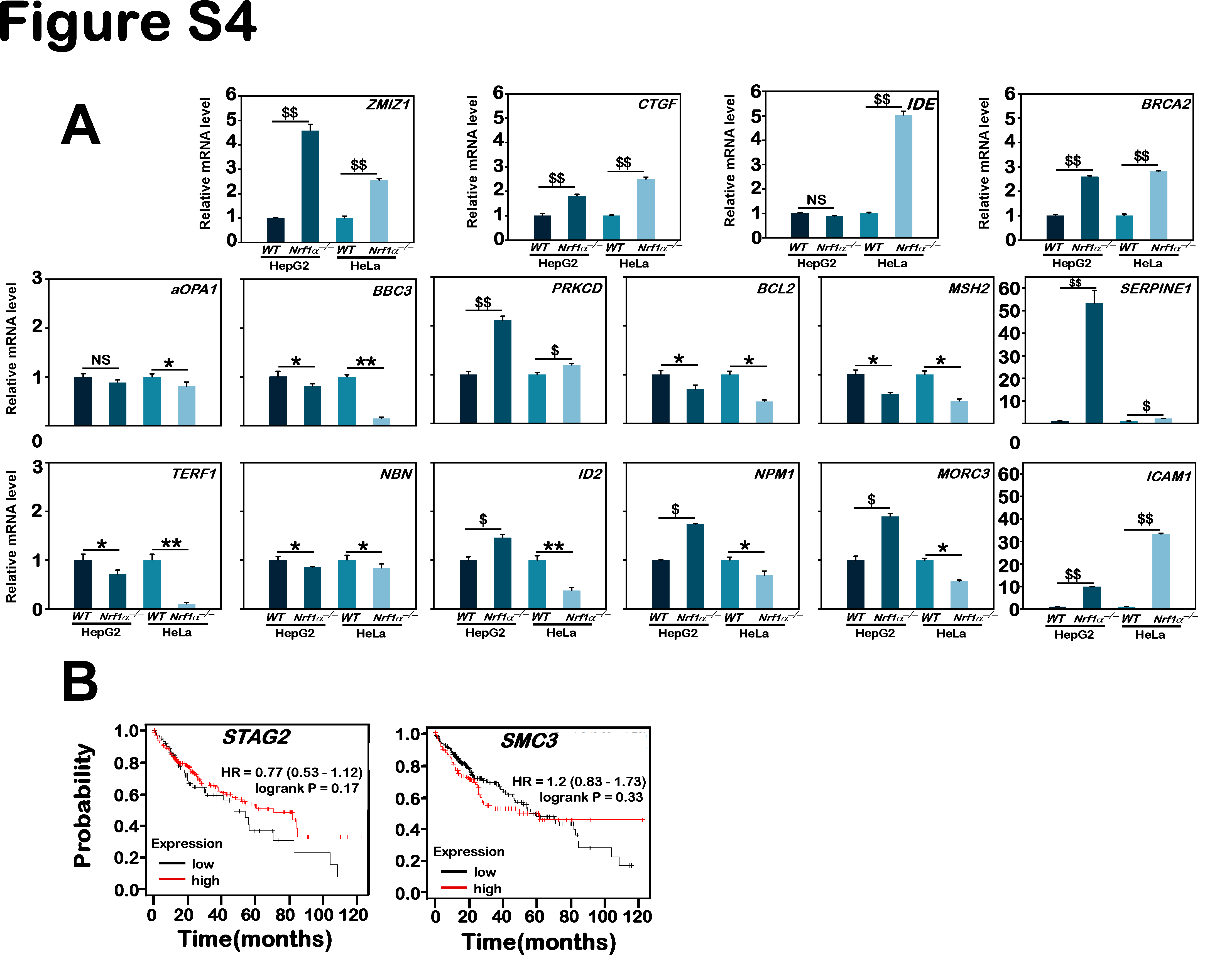

**Fig. S4. Those critical DEGs regulated by Nrf1 were further identified**

(A) Distinct mRNA expression levels of selected key genes, *aOPA1, BBC3, PRKCD, BCL2, MSH2, TERF1, NBN, ID2, NPM1, MORC3, ZMIZ1, CTGF, IDE, BRCA2, ICAM1,* and *SERPINE1* in *WT* and *Nrf1α^−/−^* cell lines were further validated by real-time qPCR. The resulting data are shown as Mean ± SD, with significant increases ($, p < 0.05; $$, p < 0.01), significant decreases (*, p < 0.05; **, p < 0.01;) or no statistical differences (NS), which were statistically determined from at least three independent experiments performed in triplicates (n = 3 × 3).

(B) The correlation analysis of STAG2 and SMC3 with relevant OS rates of patients with HCC. The data were obtained from the online Kaplan-Meier plotter database.

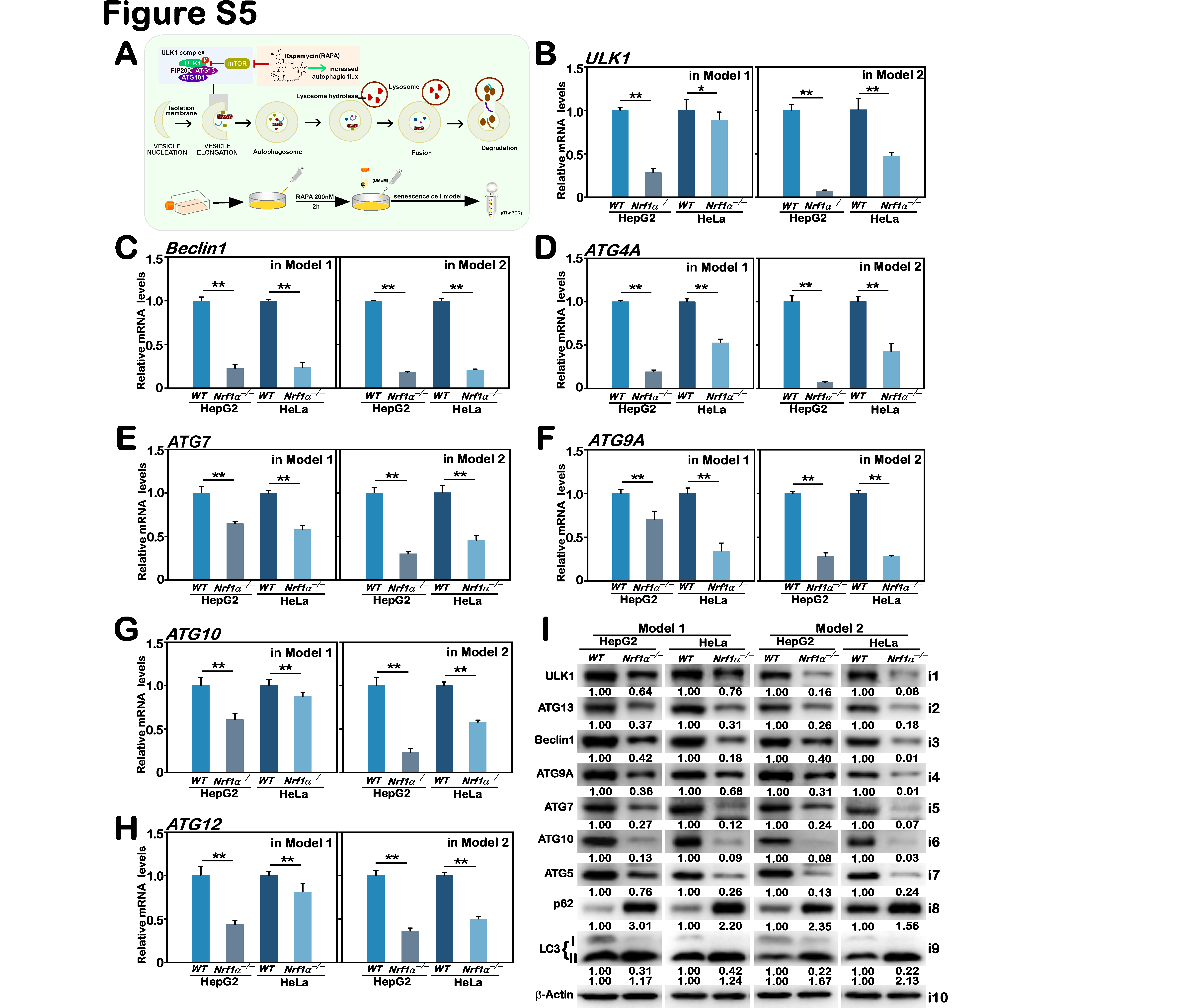

**Fig. S5. Potential effects of *Nrf1α^−/−^*-deficiency on the autophagy flux of cells treated with Rapamycin**

(A) A schematic shows changes in the flux of cell autophagy stimulated by 200 nM of Rapamycin.

(B–H) Real-time qPCR analysis of altered mRNA expression levels of autophagy-related markers, *ULK1* (B)*, Beclin1* (C)*, ATG4A*(D)*, ATG7* (E)*, ATG9A* (F)*, ATG10* (G)*,* and *ATG12* (H) in *WT* and *Nrf1α^−/−^* cell lines under distinct model senescence conditions. The resulting data are shown as Mean ± SD, with significant decreases (*, p < 0.05; **, p < 0.01), which were statistically determined from at least three independent experiments performed in triplicates (n = 3 × 3).

(I) Western blotting analysis of altered protein expression levels of ULK1, Beclin1, ATG5, ATG7, ATG9A, ATG10, ATG13, p62, and LC3 in *WT* and *Nrf1α^−/−^* cell lines under distinct model senescence conditions. The intensity of all immunoblots was quantified and then normalized by that of β-actin, before being shown as fold changes (*on the bottom* of indicated blots). The data presented herein were a representative of three independent experiments.

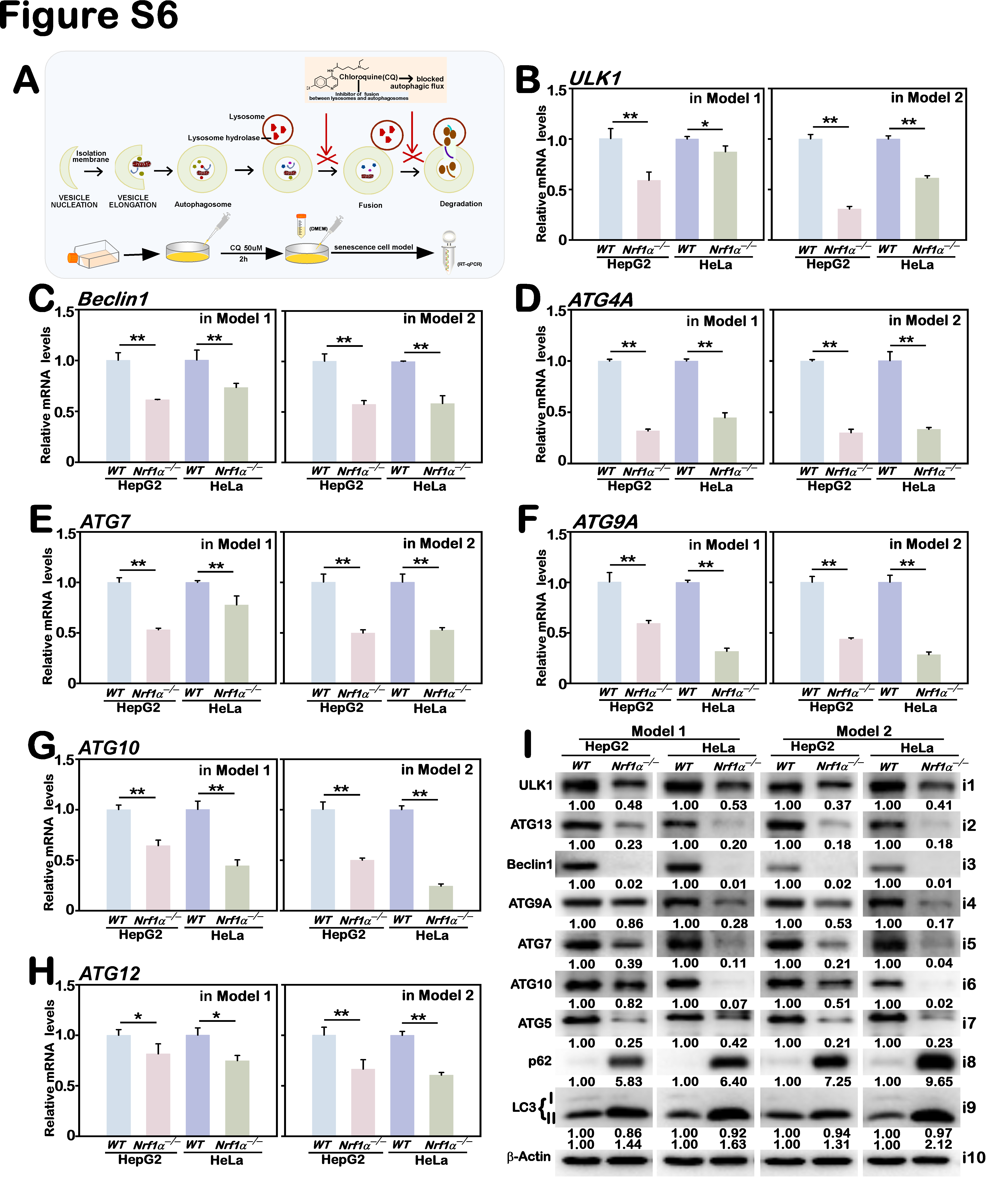

**Fig. S6. Potential effects of *Nrf1α^−/−^*-deficiency on the autophagy flux of cells treated with chloroquine**

(A) A schematic shows changes in the flux of cell autophagy stimulated by 50 µM of chloroquine.

(B–H) Real-time qPCR analysis of altered mRNA expression levels of autophagy-related markers, *ULK1* (B)*, Beclin1* (C)*, ATG4A*(D)*, ATG7* (E)*, ATG9A* (F)*, ATG10* (G)*,* and *ATG12* (H) in *WT* and *Nrf1α^−/−^* cell lines under distinct model senescence conditions. The resulting data are shown as Mean ± SD, with significant decreases (*, p < 0.05; **, p < 0.01), which were statistically determined from at least three independent experiments performed in triplicates (n = 3 × 3).

(I) Western blotting analysis of altered protein expression levels of ULK1, Beclin1, ATG5, ATG7, ATG9A, ATG10, ATG13, p62, and LC3 in *WT* and *Nrf1α^−/−^* cell lines under distinct model senescence conditions. The intensity of all immunoblots was quantified and then normalized by that of β-actin, before being shown as fold changes (*on the bottom* of indicated blots). The data presented herein were a representative of three independent experiments.

**Table S1. The key materials and relevant resources used in this work**

| Reagent or resource Source Identifier | | |
| --- | --- | --- |
| Chemicals | | |
| D-Galactose  Rapamycin  Chloroquine  Paraformaldehyde  crystalviolet solution  PMSF  Protease inhibitors  RIPA | Solarbio  Sigma-Aldrich  Solarbio  Sangon  Solarbio  Sangon  Roche  Beyotime | IG0540  553210  C9720  E672002  G1062  A100754  3271382-1  P0013C |

Antibodies

| LaminA  Ki67  p53  p21  CDK2  CyclinD  pRB  E2F1  ULK1  ATG13  Beclin1  ATG9A  ATG7  ATG10  ATG5  p62  Lc3  Keap1  Nrf1  Nrf2  STAG2  SMC3  -acti | Abcam  Wanleibio  Sangon  Sangon  ABclonal  Wanleibio  Sangon  Wanleibio  Abcam  Sangon  ABclonal  Cusabio  Abcam  Sangon  KleanAB  MBL  Abcam  ABclonal  Zhang’s(1)  Abcam  Abcam  Abcam  ZSGB-BIO | ab227176  WL01384a  D191060  D220403  A0294  WL01435a  D155057  WL02394  ab167139  D225082  A21191  CSB-PA554014  ab52472  D290115  P100534  PM045  ab192890  A17062  N/A  ab62352  ab155081  ab128919  TA-09 |
| --- | --- | --- |
| Oligonucleotides for small interference (si) RNA | | |
| Normal control FW  Normal control REV  siNrf2FW  siNrf2REV  siSTAG2FW  siSTAG2REV  siSMC3FW  siSMC3REV | Tsingke  Tsingke  Tsingke  Tsingke  Tsingke  Tsingke  Tsingke  Tsingke | UUCUCCGAACGUGUCACGUdTdT  ACGUGACACGUUCGGAGAAdTdT  CCGGCAuuuCACuAAACACAAdTdT  UuGuGuuuAGuGAAAuGCCGGdTdT  GAGAGGAAGCACUAACAGAUAdTdT  UAUCUGUUAGUGCUUCCUCUCdTdT  GUACUGGUCCUCGUGUUAUUUdTdT  AAAUAACACGAGGACCAGUACdTdT |
| Oligonucleotides for construct | | |
| STAG2-ARE1-WTFW  STAG2-ARE1-WTREV  STAG2-ARE1-MUTFW  STAG2-ARE1-MUTREV  STAG2-ARE2-WTFW  STAG2-ARE2-WTREV  STAG2-ARE2-MUTFW  STAG2-ARE2-MUTREV  STAG2-ARE3-WTFW  STAG2-ARE3-WTREV  STAG2-ARE3-MUTFW  STAG2-ARE3-MUTREV  SMC3-ARE1-WTFW  SMC3-ARE1-WTREV  SMC3-ARE1-MUTFW  SMC3-ARE1-MUTREV  SMC3-ARE2-WTFW  SMC3-ARE2-WTREV  SMC3-ARE2-MUTFW  SMC3-ARE2-MUTREV  SMC3-ARE3-WTFW  SMC3-ARE3-WTREV  SMC3-ARE3-MUTFW  SMC3-ARE3-MUTREV | Tsingke  Tsingke  Tsingke  Tsingke  Tsingke  Tsingke  Tsingke  Tsingke  Tsingke  Tsingke  Tsingke  Tsingke  Tsingke  Tsingke  Tsingke  Tsingke  Tsingke  Tsingke  Tsingke  Tsingke  Tsingke  Tsingke  Tsingke  Tsingke | cCTTGCCTGTGATCCCAGCTTACTCAGGAGGCTGAGGCAGGa  gatctCCTGCCTCAGCCTCCTGAGTAAGCTGGGATCACAGGCAAGggtac  cCTTGCCTGTGATCCCAatTTACCTAGGAGGCTGAGGCAGGa  gatctCCTGCCTCAGCCTCCTAGGTAAatTGGGATCACAGGCAAGggtac  cGAGGTTGCAGTGAGCTGAGATCGCACCACTGCACTCCAGa  gatctCTGGAGTGCAGTGGTGCGATCTCAGCTCACTGCAACCTCggtac  cGAGGTTGCAGTGAGCTAGGATCATACCACTGCACTCCAGa  gatctCTGGAGTGCAGTGGTATGATCCTAGCTCACTGCAACCTCggtac  cGCACTCCAGCTTGGGTGACAGAGCGAGAGTTTGTCTCAAa  gatctTTGAGACAAACTCTCGCTCTGTCACCCAAGCTGGAGTGCggtac  cGCACTCCAGCTTGGGTAGCAGAATGAGAGTTTGTCTCAAa  gatctTTGAGACAAACTCTCATTCTGCTACCCAAGCTGGAGTGCggtac  cCTTGAACCCAGGAGTTTGAGGCTGCAGTGAACCATGACCGAGCa  gatctGCTCGGTCATGGTTCACTGCAGCCTCAAACTCCTGGGTTCAAGggtac  cCTTGAACCCAGGAGTTTAGGGCTATAGTGAACCATAGCCGAATa  gatctATTCGGCTATGGTTCACTATAGCCCTAAACTCCTGGGTTCAAGggtac  cCAGCTCCACCCTGGGTGACAGAGCAAGATCCTGTCAAAAa  gatctTTTTGACAGGATCTTGCTCTGTCACCCAGGGTGGAGCTGggtac  cCAGCTCCACCCTGGGTAGCAGAATAAGATCCTGTCAAAAa  gatctTTTTGACAGGATCTTATTCTGCTACCCAGGGTGGAGCTGggtac  cACAAAGGGGCAGCAGTGACTCCGCACACAGAATTCCAAGa  gatctCTTGGAATTCTGTGTGCGGAGTCACTGCTGCCCCTTTGTggtac  cACAAAGGGGCAGCAGTAGCTCCATACACAGAATTCCAAGa  gatctCTTGGAATTCTGTGTATGGAGCTACTGCTGCCCCTTTGTggtac |
| Recombinant DNA | | |
| pcDNA3.1  pGL3-promoter  pRL-TK | Invitrogen  Promega  Promega | V79020  VQP0124  VQP0126 |
| Oligonucleotides for qPCR | | |
| Nrf2FW  Nrf2REV  STAG2FW  STAG2REW  SMC3FW  SMC3REW  aOPA1FW  aOPA1REW  BBC3FW  BBC3REW  PRKCDFW  PRKCDREW  BCL2FW  BCL2REW  MSH2FW  MSH2REW  TERF1FW  TERF1REW  NBNFW  NBNREW  ID2FW  ID2REW  NPM1FW  NPM1REW  MORC3FW  MORC3REW  ZMIZ1FW  ZMIZ1REW  CTGFFW  CTGFREW  IDEFW  IDEREW  BRCA2FW  BRCA2REW  ICAM1FW  ICAM1REW  SERPINE1FW  SERPINE1REW  IL1αFW  IL1αREW  IL1βFW  IL1βREW  IL6FW  IL6REW  IL8FW  IL8REW  CXCL3FW  CXCL3REW  CXCL10FW  CXCL10REW  MMP1FW  MMP1REW  ULK1FW  ULK1REW  Beclin1FW  Beclin1REW  ATG4AFW  ATG4REW  ATG7FW  ATG7REW  ATG9AFW  ATG9AREW  ATG10FW  ATG10REW  ATG12FW  ATG12REW  β-actin FW  β-actin REV | Tsingke  Tsingke  Tsingke  Tsingke  Tsingke  Tsingke  Tsingke  Tsingke  Tsingke  Tsingke  Tsingke  Tsingke  Tsingke  Tsingke  Tsingke  Tsingke  Tsingke  Tsingke  Tsingke  Tsingke  Tsingke  Tsingke  Tsingke  Tsingke  Tsingke  Tsingke  Tsingke  Tsingke  Tsingke  Tsingke  Tsingke  Tsingke  Tsingke  Tsingke  Tsingke  Tsingke  Tsingke  Tsingke  Tsingke  Tsingke  Tsingke  Tsingke  Tsingke  Tsingke  Tsingke  Tsingke  Tsingke  Tsingke  Tsingke  Tsingke  Tsingke  Tsingke  Tsingke  Tsingke  Tsingke  Tsingke  Tsingke  Tsingke  Tsingke  Tsingke  Tsingke  Tsingke  Tsingke  Tsingke  Tsingke  Tsingke  Tsingke  Tsingke | TCAGCGACGGAAAGAGTATGA  CCACTGGTTTCTGACTGGATGT  GAGCCACTTAGTGGAGAGGAAG  GTAAGCACCCTTTTTCCTGTCCC  ATGCGTGGAAGTCACTGCTGGA  GGCAGAAAAGTAACCTCTCCAGG  GTGGTTGGAGATCAGAGTGCTG  GAGGACCTTCACTCAGAGTCAC  ACGACCTCAACGCACAGTACGA  CCTAATTGGGCTCCATCTCGGG  AACCATGAGTTTATCGCCACC  AGCGTTACATTGCCTGCATTT  ATCGCCCTGTGGATGACTGAGT  GCCAGGAGAAATCAAACAGAGGC  CAGCAGTCAGAGCCCTTAACCT  GAGAGGCTGCTTAATCCACTGG  CATGGAACCCAGCAACAAGACC  CTGCTTTCAGTGGCTCTTCTGC  AGGTGGGGAAGCTAGGTTGAT  CACCGCCAATCCAATTTCTGC  TTGTCAGCCTGCATCACCAGAG  AGCCACACAGTGCTTTGCTGTC  GCCAGTGCATATTAGTGGACAGC  GGAACCTTGCTACCACCTCCAG  TCTTCTCCTAAGGAAAGTGTTCCA  GAGGAACGAGTAGAAAGTCTCCG  TGTCCTCCTGGTGCGAAGA  CCCATCACTGTGCGACAGA  CTTGCGAAGCTGACCTGGAAGA  CCGTCGGTACATACTCCACAGA  TACCTCCGCTTGCTGATGACTG  ACAGGAGCTGAGGTATGAAGGC  GGCTTCAAAAAGCACTCCAGATG  GGATTCTGTATCTCTTGACGTTCC  GTATGAACTGAGCAATGTGCAAG  GTTCCACCCGTTCTGGAGTC  CTCATCAGCCACTGGAAAGGCA  GACTCGTGAAGTCAGCCTGAAAC  TGTATGTGACTGCCCAAGATGAAG  AGAGGAGGTTGGTCTCACTACC  CCACAGACCTTCCAGGAGAATG  GTGCAGTTCAGTGATCGTACAGG  AGACAGCCACTCACCTCTTCAG  TTCTGCCAGTGCCTCTTTGCTG  GAGAGTGATTGAGAGTGGACCAC  CACAACCCTCTGCACCCAGTTT  TTCACCTCAAGAACATCCAAAGTG  TTCTTCCCATTCTTGAGTGTGGC  GGTGAGAAGAGATGTCTGAATCC  GTCCATCCTTGGAAGCACTGCA  ATGAAGCAGCCCAGATGTGGAG  TGGTCCACATCTGCTCTTGGCA  GCAAGGACTCTTCCTGTGACAC  CCACTGCACATCAGGCTGTCTG  CTGGACACTCAGCTCAACGTCA  CTCTAGTGCCAGCTCCTTTAGC  CCAAGCCAGAAGTGACAACCAC  GACAGACCTTCAAGTTGAGTTCC  CGTTGCCCACAGCATCATCTTC  CACTGAGGTTCACCATCCTTGG  GCTTCCTCAAGGAGCAGGTTCA  CCACATTTGCGATAAGGCTCAGG  GGTGATAGTTGGGAATGGAGACC  GTCTGTCCATGGGTAGATGCTC  GGGAAGGACTTACGGATGTCTC  AGGAGTGTCTCCCACAGCCTTT  CATGTACGTTGCTATCCAGGC  CTCCTTAATGTCACGCACGAT |
| Software and Algorithms | | |
| Canvas 9  Origin  Excel  FlowJo 7.6.1  SPSS 22.0  Cytoscape  Premier 5 | Cancas GFX, Inc.  OriginLab  Microsoft  FlowJo  IBM  (2)  PREMIER Biosoft International | https://www.canvasgfx.com/  <https://www.originlab.com/>  https://www.microsoft.com/  https://www.flowjo.com/  <https://www.ibm.com/>  [http://www.cytoscape.org/](http://www.cytoscape.org/" \t "https://www.mdpi.com/2072-6694/10/12/_blank)  [https://www.PremierBiosoft.com/](https://www.premierbiosoft.com/" \t "https://www.mdpi.com/2072-6694/10/12/_blank) |
| [1] Zhang, Y.G.; Hayes, J.D. Identification of topological determinants in the N-terminal domain of transcription factor Nrf1 that control its orientation in the endoplasmic reticulum membrane. *Biochem. J.* **2010**, *430*, 497–510.  [2] Shannon, P.; Markiel, A.; Ozier, O.; Baliga, N.S.; Wang, J.T.; Ramage, D.; Amin, N.; Schwikowski, B.; Ideker, T. Cytoscape: A software environment for integrated models of biomolecular interaction networks. *Genome Res.* **2003**, *13*, 2498–2504. | | |
